# Gene flow erodes core gut microbiota, but not microbiome function, across a natural primate hybrid zone

**DOI:** 10.64898/2025.12.10.693535

**Authors:** Mariah E. Donohue, Jaesylin Stephens, David R. Weisenbeck, Katherine R. Martin, Jeffrey H. Frederick, Andrew N. Black, William J. Ranaivosolo, Jean Patrique Raheimanjato, Mai Fahmy, Michael Bliss, Patricia C. Wright, David W. Weisrock, Kathryn M. Everson

**Affiliations:** Dept. of Biology, University of Kentucky, Lexington, KY, 40508; Dept. of Biological Sciences, SUNY Binghamton, Binghamton, NY, 13902; Dept. of Integrative Biology, Oregon State University, Corvallis, OR, 97330; Center for Quantitative Life Sciences, Oregon State University, Corvallis, OR, 97330; Centre ValBio Research Station, Ranomafana, Madagascar; Dept. of Anthropology, University of Antananarivo, Antananarivo, Madagascar; Dept. of Biology, Fordham University, Bronx, NY, 10458; Ecology & Evolutionary Biology Interdisciplinary Program, Texas A&M University, College Station, TX, 77843; Dept. of Ecology & Conservation Biology, Texas A&M University, College Station, TX, 77843; Dept. of Anthropology, Stony Brook University, Stony Brook, NY, 11794

**Keywords:** Hybridization, primates, gut microbiome, population genomics, phylosymbiosis

## Abstract

The gut microbiome is shaped by host ecology and evolution, but the relative influences of each remain unresolved in natural hybrid zones where species boundaries are permeable. We investigated these dynamics using a hybrid zone formed between two species of brown lemur in southeastern Madagascar, *Eulemur rufifrons* and *E. cinereiceps.* The hybrids occupy a novel ecological niche and were previously proposed as a hybrid species, though this had never been tested with genome-wide data. Here, we combined low-coverage host genomes derived from fecal DNA with 16S rRNA gut microbiome profiles across a 180-kilometer transect. Population genomic analyses recovered two population clusters connected by a hybrid zone biased towards *E. rufifrons* ancestry. Though our data did not support a hypothesis of hybrid speciation, the hybrid zone was dominated by later-generation crosses. Microbiome beta diversity was significantly correlated with host genomic distance after accounting for geography and ecology. Hybrid microbiomes were distinguishable from parentals, but this divergence was not driven by the accumulation of novel microbiota. Instead, hybrids lost core microbes present in both parental species. Despite this loss of diversity, hybrid gut microbiome function was nearly indistinguishable from parentals. Together, this suggests microbiome adaptation is not a necessary pre-requisite for host ecological adaptation. We also demonstrate that species-specificity of the gut microbiome can emerge in the face of interspecific gene flow, providing new insight into host–microbiome dynamics during the early stages of speciation.

## INTRODUCTION

The study of evolution has expanded beyond the host itself to include diverse microbial communities that live in and on every plant and animal [1]. These symbiotic microbes are integral to host health, survival, and ecological adaptation [2, 3, 4]. Thus, understanding the causes and consequences of microbiome variation offers important insights into how host species evolve as multi-species systems, collectively termed holobionts [5]. Host ecological factors such as habitat [6, 7, 8, 9] and diet [10, 11, 12, 13] are known to powerfully and predictably shape the microbiome. However, the effects of evolutionary factors, such as hybridization, population divergence, and speciation, remain poorly understood.

One of the strongest lines of evidence for an evolutionary influence on microbiome assembly is phylosymbiosis, a pattern in which microbiome dissimilarity increases with phylogenetic distance [14]. Phylosymbiosis has been observed across the Tree of Life, and can arise via three primary mechanisms: ecological filtering by phylogenetically conserved host traits [15, 16, 17, 18], dispersal limitation that restricts microbial exchange among host species [19,20], and reciprocal host–microbiome coevolution [21, 22, 23]. Intraspecific variation rarely exceeds interspecific variation, making the gut microbiome a diagnostic character of many host species, particularly mammals [24, 20].

Our understanding of co-divergence is biased towards “good” host species with complete reproductive isolation. However, reproductive barriers rarely evolve instantaneously [25]. Instead, varying degrees of interspecific gene flow characterize early stages of speciation [26, 27, 28]. This can have a range of genomic consequences, from the introgression of beneficial alleles [29, 30, 31] to the formation of new hybrid species that are reproductively isolated from both parental lineages [32, 33], or alternatively the reinforcement of species boundaries due to infertile F1 hybrids [34, 35].

Likewise, hybridization has many potential outcomes in the gut microbiome. In some cases, hybrid microbiomes are intermediate, representing a blend of both parental species [36, 37, 38]. Intermediate microbiomes increase the risk of mismatches analogous to Dobzhansky-Muller incompatibilities [39], but may also preserve, or even enhance, the functional diversity of parental microbiomes in hybrid offspring. Alternatively, hybrid microbiomes may be transgressive, meaning they bear little resemblance to either parent and can foster major phenotypic divergence [40]. Where transgressive microbiomes are advantageous, they can enable hybrids to adapt to novel ecological conditions relative to parental species [41, 39]. However, many studies have found that transgressive microbiomes yield negative host outcomes, as exemplified by *Nasonia* wasps [42] and house mice [43].

Hybridization is increasingly recognized as a feature of adaptive radiations [44, 45, 46, 47]. This is especially well documented in lemurs, which account for nearly 20% of extant primate diversity despite endemism on an island encompassing < 1% of Earth’s land area. Lemurs split from other Strepsirrhines ∼50 MYA and have since diversified into five families, 15 genera, and > 110 species. Due to multiple independent bursts of speciation, species diversity is heterogeneous across the Lemur Tree. The genus Eulemur is among the most explosive of its clades, experiencing an increased diversification rate relative to other strepsirrhine clades of a similar age [46]. Phylogenetic analyses support a deep and rampant history of hybridization in Eulemur [47, 48, 49], with at least five active Eulemur hybrid zones across Madagascar [48].

Hybrids of *Eulemur rufifrons* and *E. cinereiceps* are of particular interest, as they have been proposed as a putative hybrid species [50] based on two primary lines of evidence. First, the hybrids possessed microsatellite alleles absent in parental species, suggesting assortative mating and some degree of reproductive isolation [51, 52]. Second, hybrids occupy a distinct ecological niche compared to both parents. They live in transitional rainforests with unique climatic trends throughout the year [52, 53] and exhibit extreme specialization on a diet of fruit [50]. Despite evidence of evolutionary and ecological divergence, formal tests of this hypothesized hybrid speciation event have never been performed. Nonetheless, this population offers a compelling natural system for understanding microbiome assembly in ecologically successful hybrids.

Here, we apply a holobiont framework to characterize the evolutionary history of *E. rufifrons x E. cinereiceps*. With fecal-derived genomic DNA, we test a hypothesis of hybrid speciation by evaluating population structure across a transect encompassing the hybrid zone and adjacent parental populations. We then use this information to elucidate the relative effects of host ecology and evolutionary history on gut microbiome composition, diversity, and predicted function. These results enhance our understanding of host-microbiome co-evolution given complex histories of interspecific gene flow, while also providing new insight into the biology of endangered primate species.

## METHODS

### Ethics statement

All the research complies with applicable laws on sampling from natural populations and animal experimentation (including the ARRIVE guidelines). Fecal collection protocols were approved by the University of Kentucky IACUC (2021-3814) and Madagascar National Parks (132/22/MEDD/SG/DGGE/DAPRNE/SCBE.Re). This research adhered to the American Society of Primatologists Principles for the Ethical Treatment of Non-Human Primates and all legal requirements in Madagascar.

### Sample collection

We collected fecal samples along a 180 km north-south transect extending from *E. rufifrons* populations in the north, through the entire purported hybrid region, and into nearly all known *E. cinereiceps* populations in the south (Fig. 1A). The transect fell mostly within the Corridor Forestier d’Ambositra Vondrozo (COFAV), a humid-forest corridor encompassing two national parks (Ranomafana and Andringitra) and a narrow tract of montane rainforest connecting them to each other and more distant protected areas [54]. We also sampled isolated populations of *E. cinereiceps* in Manombo Special Reserve. At each site, we aimed to collect fecal samples from all individuals in at least two groups. Samples were collected within five minutes of deposition using sterilized tweezers, preserved in 96% ethanol, and kept at room temperature until transport to the University of Kentucky, where they were then transferred to a -20°C freezer.

**Fig. 1:**
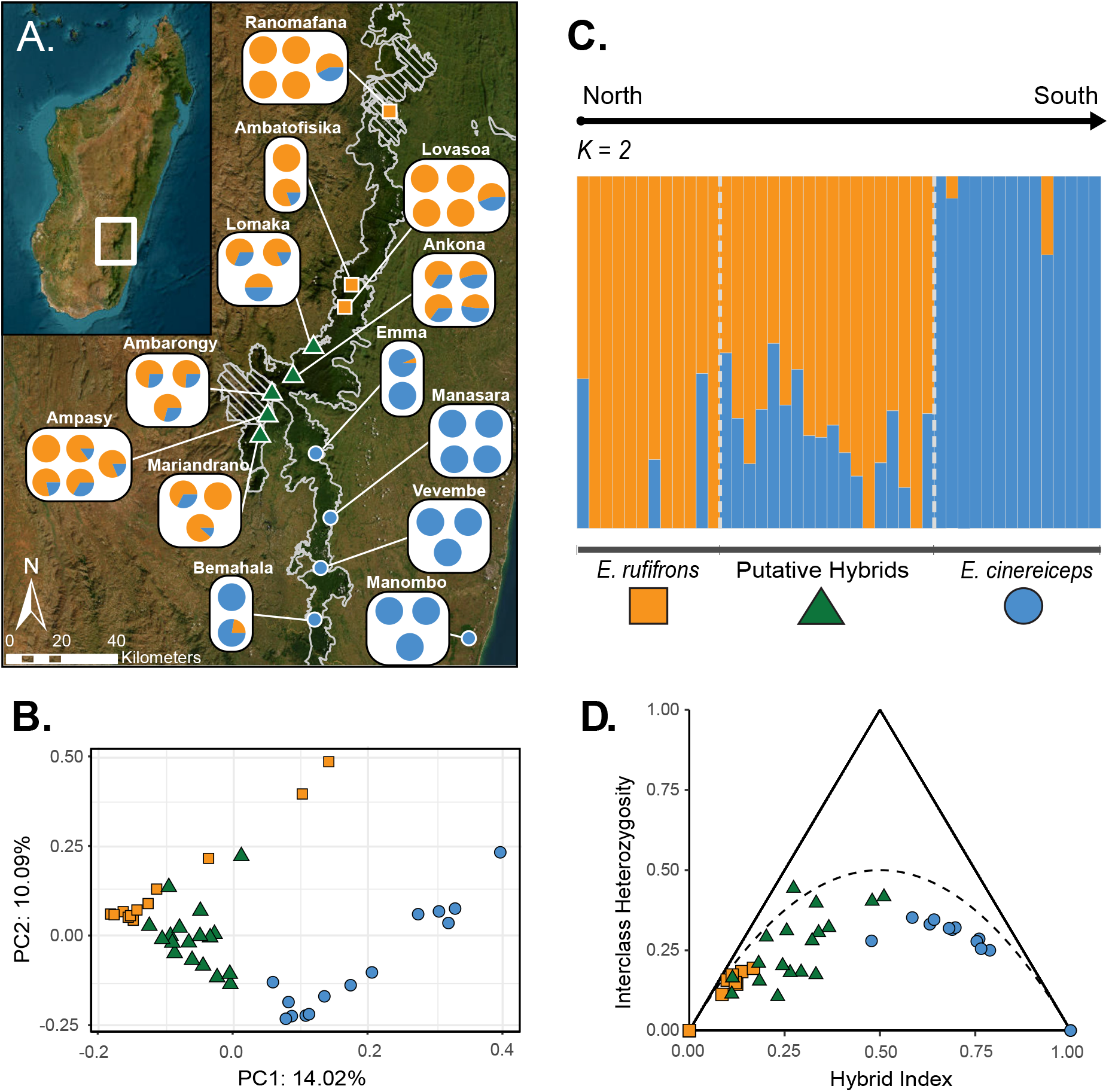
Population genomic analyses of the *E. cinereiceps* x *E. rufifrons* hybrid zone. In each panel, orange color denotes *E. rufifrons*, green hybrid, and blue *E. cinereiceps*. **(A)** Map of sampling sites and admixture results. Hatched regions denote national parks, with Ranomafana in the north and Andringitra in the south; white outlined regions represent the COFAV. Points on the map illustrate *a priori* population assignments based on previous genetic studies (Delmore et al., 2013). Pie charts show proportions of genetic ancestry for each individual sequenced in this study based on *K = 2*, the best supported model (Fig. S1). **(B)** PCA of population structure. Hybrids appear intermediate to parentals, with some overlap between hybrids and *E. rufifrons*, and clear separation of *E. cinereiceps.* **(C)** Bar charts of admixture results at *K = 2* along a north to south continuum, with *a priori* population assignments imposed along the x-axis. **(D)** Triangle plot showing the hybrid index (x-axis) and interclass heterozygosity (y-axis) of every individual in the dataset. Additional triangle plots with varying allele frequency difference thresholds are shown in Fig. S4.

From these samples, we generated low-coverage whole-genome sequence data from 44 individual lemurs (putative identifications: 14 *E. cinereiceps*, 12 *E. rufifrons,* 18 hybrids; Table S1), which were then used to generate a dataset of single nucleotide polymorphisms (SNPs) for population genomic and demographic analyses. We also generated 16S rRNA gut microbiome profiles for 105 fecal samples (putative identifications: 38 *E. cinereiceps*, 24 *E. rufifrons,* and 41 hybrids; Table S2).

### Lemur genomic data generation

We extracted DNA from ∼0.2 g of feces using the QIAGEN QIAamp® Fast DNA Stool Mini Kit following the “Isolation of DNA from Stool for Human DNA Analysis” manufacturer protocol. We then preferentially enriched eukaryotic DNA using the FecalSeq protocol (supplemental methods) [55]. Whole-genome libraries were prepared at the Florida State University Molecular Cloning Lab using the NEBNext® Ultra™ II DNA Library Prep Kit for Illumina®. Libraries were sequenced on a NovaSeq X Series Illumina sequencer to generate 150 bp paired-end reads at a sequencing depth of 30,000,000 reads per sample.

### Gut microbiome data generation

We extracted DNA from ∼0.2 g of feces using the Qiagen QIAamp® PowerFecal® Kit following the manufacturer protocol. The V4 region of 16S rRNA was PCR amplified using dual-indexed primers (supplemental methods) [56]. Pooled libraries were sequenced on an Illumna MiSeq at the University of Kentucky Genomics Core.

### Data analysis: Lemur population genomics

After bioinformatic processing of the lemur genomic dataset (supplemental methods), we estimated F_ST_ between *E. cinereiceps*, *E. rufifrons*, and putative hybrids in angsd [57] based on site frequency spectra (SFS). We calculated the two-dimensional SFS between each taxonomic unit in a pairwise fashion using the “realSFS” command. The resulting two-dimensional SFSs for each pair were used in the realSFS “fst index” command as priors to calculate binary F_ST_ files. From the binary F_ST_ files, we extracted the F_ST_ values using the realSFS “fst stats” command.

We performed a PCA with pcangsd v.0.9757 [58] on the covariance matrix calculated from the genotype likelihoods. We visualized the PCA using ggplot v. 3.5.2 [59]. To infer the number of genetic clusters in our dataset (K) and the admixture proportions of each sample, we implemented a model-based approach with ngsadmix v. 2 [60]. Genotype likelihoods were used as input for 100 runs of ngsadmix, each assessing K =1 through K = 10. For each run, we specified 50,000 expectation-maximization algorithm iterations (-maxiter), a tolerance for convergence of 1e-9 (-tol) and a log likelihood difference in 50 iterations of 1e-9 (-tolLike50). We used the clumpak webserver [61] to determine the most likely K from the log likelihood scores of each ngsadmix run [62], and then clumpp v.1.12 [63] to aggregate individual runs into merged Q-matrices. We visualized the merged Q-matrices using pophelper v.2.3.1 [64].

We estimated triangle plots via triangulaR [65] using the VCF file with called genotypes. When identifying diagnostic SNPs (function “alleleFreqDiff”) we treated the northernmost *E. rufifrons* population (Ranomafana) as parental species A and the southernmost *E. cinereiceps* population (Manombo) as parental species B. Because parental species harbored few fixed differences, we explored a range of allele frequency difference thresholds ranging from 0.5 to 1.0. We calculated hybrid indices using the “hybridIndex” function and generated triangle plots using the function “triangle.plot” with default settings.

We evaluated landscape-level patterns of population genomic variation using a series of spatially explicit tests of IBD and IBE [66]. We performed mixed matrix regression with randomization (MMRR) [67] and generalized dissimilarity modeling (GDM) [68] comparisons between combinations of population pairs (*E. rufifrons* + hybrids and *E. cinereiceps +* hybrids), plus a third test treating all samples as a single species. MMRR and GDM analyses used the same suite of input data: coordinate data for each sample, BioClimatic data from WorldClim2 (http://www.worldclim.org) [69], and matrices of genetic distance generated from SNP data (supplemental methods). Analyses were conducted in R using a derivation of the Algatr pipeline (supplemental methods) [70].

### Bioinformatics and data analysis: Gut microbiome data

From the microbial 16S rRNA dataset, we removed samples with fewer than 1000 reads, retaining 95 total (putative identifications: 37 *E. cinereiceps*, 24 *E. rufifrons*, 36 hybrids). All bioinformatic processing and diversity analyses were conducted in QIIME2 v2024.10.1 [71]. Reads were filtered, denoised, and chimera-removed with the DADA2 pipeline [72]; sequences with < 20 reads and those matching mitochondria, Eukaryota, and Archaeplastida were removed. Alpha and beta diversity were calculated using “core_diversity_analyses.py,” rarefying at 5,943 reads/sample. The rarefied table was also used to compute relative abundance via the “qiime taxa barplot” function. We visualized the top five most abundant phyla per sample with a stacked barplot using ggplot2.

We used QIIME2’s Random Forest classifier to predict host species identity from microbial composition, splitting the data into a training set (80%) and a test set (20%) and running 10,000 simulations. Differences in alpha diversity (observed amplicon sequence variants, or ASVs) were tested with Kruskal-Wallis [73], and homogeneity of variance was assessed with the Fligner-Killeen test [74]. Results were visualized with ggplot2 violin plots. To examine variation in beta diversity, we first assessed the degree of microbiome divergence between parentals and hybrids using pairwise permutational multivariate analysis of variance (PERMANOVA) with multiple-comparison corrections and visualized patterns using qiime2r Principal Coordinate Analysis (PCoA) plots (https://github.com/jbisanz/qiime2R). To compare the relative effects of ecology and evolutionary history on beta diversity, we ran an adonis test setting beta diversity as the dependent variable and the formula “Species+Site/Group+Sex” as the independent variable. Here, host “species” (classification as *E. rufifrons, E. cinereiceps,* or hybrid) represents evolutionary history and “site” (sampling location) serves as a proxy for ecology. “Group”, which refers to an individual’s social group, was set as a nested factor within “site.” We also used Mantel and Partial Mantel tests to evaluate the relative effects of geography, ecology, and genomic ancestry on microbiome beta diversity, using the same input matrices that were used in the host MMRR and GDM analyses above (i.e., geographic distance, ecological distance calculated using bioclimatic variables, and genetic distance). In this analysis, whenever possible we used the same individual for both gut microbiome and host genomic data. When the same individual was not available, we used the next best replacement (an individual of the same sex from the same social group). Results were visualized with scatter plots using ggplot2.

We predicted microbial functional potential using KEGG orthologs [75] generated with “picrust2_pipeline.py” [76]. Functional divergence was visualized with ggpicrust’s “pathway_pca” [77]. We also converted KO abundance to a distance matrix, and tested host species’ effects on predicted function using “adonis2” in vegan [78]. We then identified KOs differing in abundance between hybrids and parentals using MaAsLin 2 with FDR correction [79]. The KOs were functionally annotated using the BRITE hierarchy [80]. As an additional test of predicted microbiome function, we used relative abundance data computed as the Firmicutes/Bacteroidetes (F/B) ratio for each host taxon. Firmicutes is typically associated with the consumption of fibrous foods, while Bacteroidetes increases with greater sugar consumption [81,82]. Differences in the F/B ratio were assessed using a Kruskal-Wallis test.

To quantify the degree of microbial novelty in hybrids relative to parents, we computed 4H indices using the R package “HybridMicrobiomes” [39]. This analysis uses core microbiomes (the specific set of ASVs consistently shared across all samples) of each host class (here, E. rufifrons, E. cinereiceps, and hybrids) to compare the relative fit of four conceptual models summarizing how hybrid microbiomes may differ from parentals. In the Union Model, hybrids host all of the microbial taxa in at least one parent, but harbor no unique, hybrid-specific taxa. In the Intersection Model, hybrids host only those microbial taxa common to both parents. In the Gain Model, hybrids exclusively host novel microbial taxa not found in either parent. And in the Loss Model, hybrids are missing microbial taxa found in one or both parents. Importantly, these models are not mutually exclusive, and data are expected to fall along a spectrum spanning completely uniform and completely divergent microbiomes [39]. Analyses were performed using the “FourHbootstrap” function with equalized sample sizes among host categories. The number of individuals sampled per group during each bootstrap replicate was set to the size of the smallest host category. One hundred bootstrap replicates were performed for each analysis. To evaluate sensitivity to core microbiome definitions, analyses were repeated across ten core thresholds ranging from 10% to 100% prevalence. For each threshold, the intersection, union, gain, and loss indices were calculated from the bootstrap centroid estimates. 4H metrics across core thresholds were visualized using line plots generated with ggplot2.

## RESULTS

### Putative hybrids are genetically intermediate to both parentals

Population structure analyses indicated widespread admixture across the transect. In a scenario with two population clusters (*K* = 2), the best supported model (Fig. S1), *E. cinereiceps* had lower levels of admixture (2/14 individuals were admixed) than *E. rufifrons* (3/12 individuals) and all but two samples recovered from the purported hybrid zone were admixed (Fig. 1B). Weighted F_ST_ was lowest between hybrids and *E. rufifrons* (0.038 compared to 0.072 between hybrids and *E. cinereiceps*; Table S3). Principal Component Analysis (PCA) showed overlap between hybrid and *E. rufifrons* clusters, forming a distinct cluster along PC1 (Fig. 1C).

Triangle plots, which assign individuals to hybrid classes (e.g., F1s, F2s, or backcrosses) [65], also supported distinct clusters for *E. rufifrons* and *E. cinereiceps* (Fig. 1D; S2). Hybrids had intermediate hybrid index values, but interclass heterozygosity never exceeded 0.5, as would be expected if our dataset included F1 individuals.

Landscape genomic analyses identified weak isolation-by-distance (IBD) between *E. rufifrons* and putative hybrids, isolation-by-environment (IBE) between *E. cinereiceps* and putative hybrids, and both IBD and IBE when all samples were treated as single deme (Table S4, S5; extended results in supplemental materials).

### Hybrid gut microbiomes are intermediate relative to parental species

After filtering, we recovered a total of 7,448 ASVs (mean frequency = 644; range = 2- 300,380). Across all samples, Bacteroidetes and Firmicutes were the most dominant microbial phyla, together accounting for 46% ± 11.2 and 33% ± 8.3 of total microbial diversity, respectively. The F/B ratio did not differ across host species (p > 0.05) (Fig. S2). The remaining ∼20% of microbial abundance was dominated by Actinobacteria (5% ± 2.9), Proteobacteria (4% ± 3.9), Cyanobacteria (3.6% ± 3.2), and Synergistota (2.6% ± 2.2), with little variation between hybrids and parentals (Fig. S3).

Using 16S rRNA gut microbiome profiles from samples across the hybrid transect, we found that the Random Forest algorithm accurately classified host identity 85% of the time, with the greatest source of model uncertainty arising in differentiating hybrids from *E. rufifrons* (Fig. 2A). Alpha diversity did not differ between host species (Kruskal-Wallis *H-value* = 1.38; *p-value* = 0.50; Fig. 2B), though hybrids exhibited higher variance than parentals (χ² = 5.98; *p- value* = 0.05). Comparisons of 4H models (Intersection, Union, Gain, and Loss) across core microbiome thresholds identified Intersection and Loss as the best fit to our data. The Intersection Model dominated across most thresholds, but the importance of Loss increased as core thresholds became more stringent. In contrast, Gain and Union remained comparatively low across analyses (Fig. 2C).

**Fig. 2:**
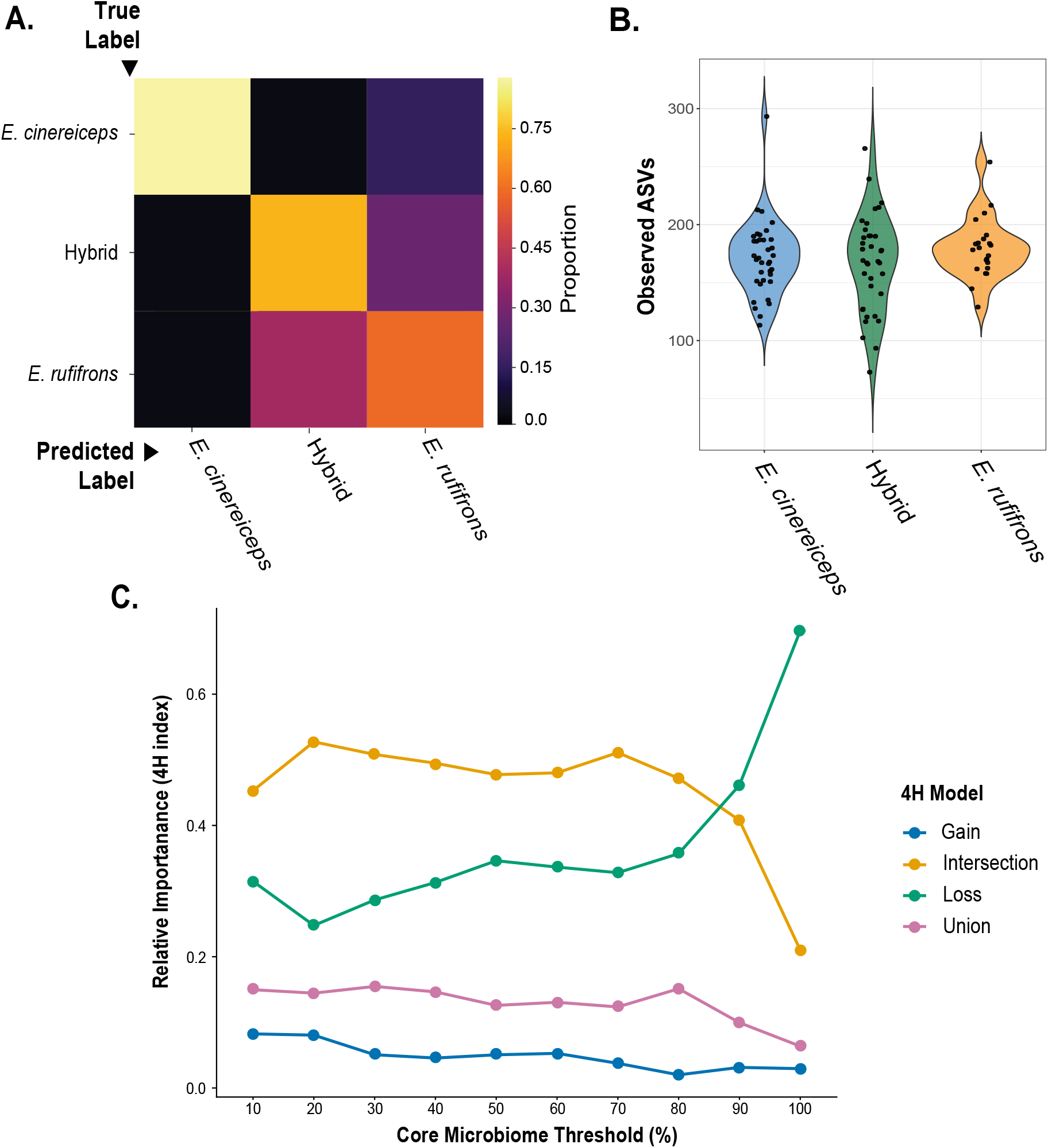
Comparison of gut microbiome composition and alpha diversity between hybrids and parentals. **(A)** Heatmap demonstrating the accuracy of QIIME2’s machine learning model in assigning host species identification based on gut microbiome composition. Samples belonging to *E. cinereiceps* were seldom misidentified, whereas more substantial model confusion arose discerning *E. rufifrons* and hybrids. **(B)** Violin plot of alpha diversity. The number of observed ASVs did not differ significantly across host species. Variance also did not differ significantly (Fligner-Killeen p-value = 0.05), though hybrid alpha diversity dispersion marginally exceeds parentals. **(C)** Changes in the relative contribution of the four hybrid microbiome assembly models across core microbiome thresholds ranging from 10% to 100%. The 4H analysis partitions the hybrid gut microbiome into four components: intersection (microbial taxa shared between both parental species), gain (novel taxa unique to hybrids), loss (parental taxa absent from hybrids), and union (taxa present in either parental species but not shared). Values represent the centroid of 100 bootstrap replicates at each core microbiome threshold, with each bootstrap subsampled to the smallest host group size. Across most thresholds, the intersection component accounted for the largest proportion of the hybrid microbiome, whereas the gain and union components remained consistently low. The loss component increased with more stringent core thresholds.

Beta diversity exhibited clear signals of divergence among host species, as Principal Coordinates Analysis (PCoA) of Jaccard beta diversity depicted three clusters with a primary separation along PCoA1; parental microbiomes were largely non-overlapping and hybrids were intermediate, but closer to *E. rufifrons* (Fig. 3A). Pairwise permutational multivariate analysis of variance (PERMANOVA) validated this visual observation, revealing high dissimilarity between *E. rufifrons* and *E. cinereiceps* (*Pseudo-F* = 4.32) and hybrids and *E. cinereiceps* (*Pseudo-F* = 3.29), and low dissimilarity between *E. rufifrons* and hybrids (*Pseudo-F* = 1.87). Adonis tests showed the effects of site (*R^2^* = 0.19) and group identity (*R^2^* = 0.12) were stronger than species (*R^2^* = 0.08) (Table S6). However, Mantel and Partial Mantel tests found genomic distance to be the best predictor of gut microbiome distance. Using Mantel tests, Jaccard beta diversity was significantly correlated with genomic distance (*R^2^*= 0.26; p = 0.013), geographic distance (*R^2^* = 0.24; p = 0.006), and environmental distance (*R^2^*= 0.19; p = 0.043) (Table S7). After accounting for geography and environment, genomic distance remained a significant predictor of beta diversity (Table S8). These trends were echoed across Bray-Curtis, Weighted UniFrac, and Unweighted UniFrac beta diversity metrics (Table S7, S8).

**Fig. 3:**
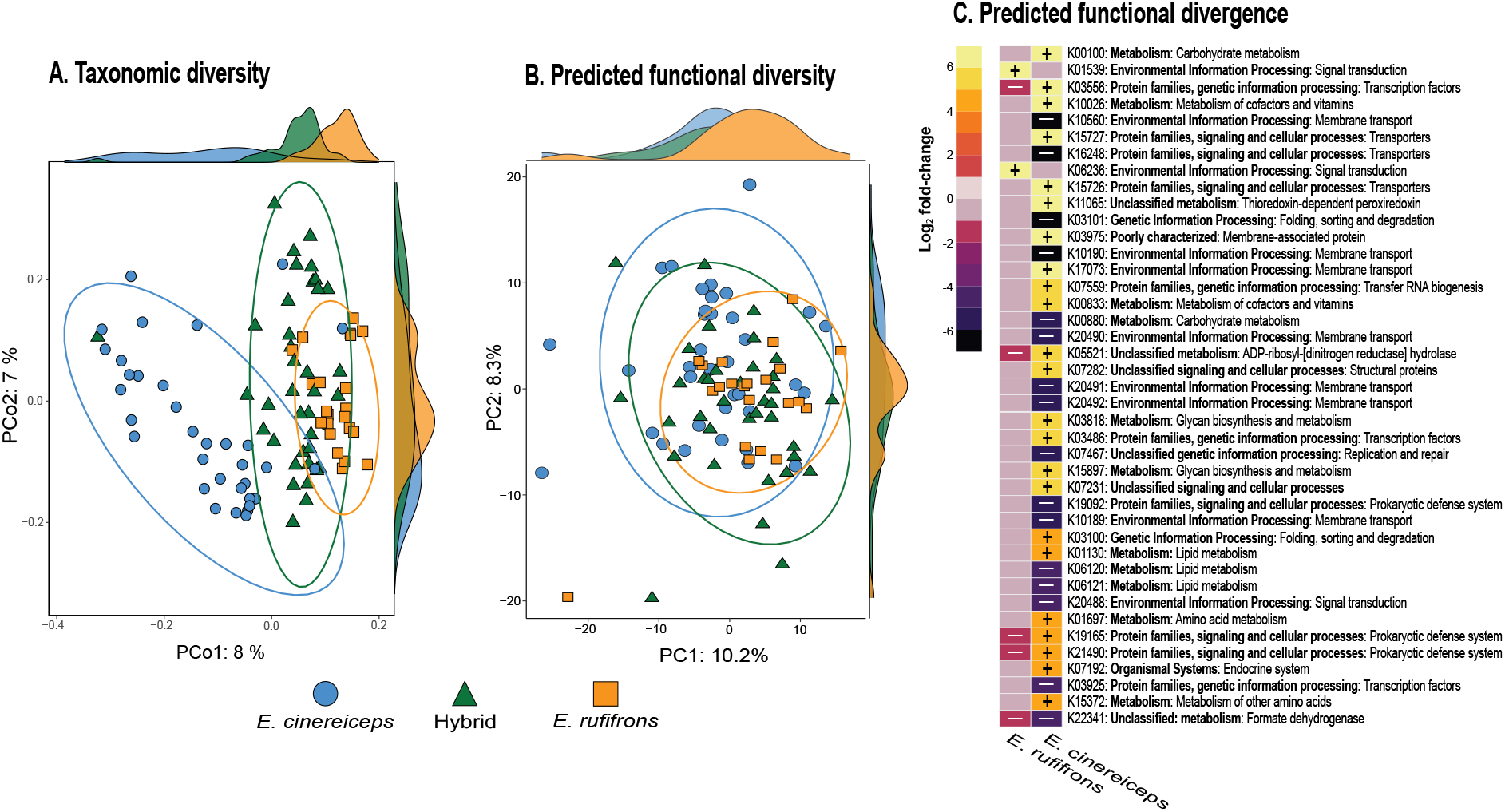
Comparison of gut microbiome beta diversity and predicted function between hybrids and parentals. **(A)** Principal Coordinates Analysis (PCoA) of Jaccard beta diversity using microbial taxonomy data, with colors and shapes corresponding to host species identification. Ellipses represent 95% confidence intervals; density plots illustrate sample distribution along the first and second axes. Hybrid samples appear intermediate to parentals, with clear separation of *E. cinereiceps* from an overlapping cluster of both hybrid and *E. rufifrons* samples on PCo1. **(B)** Principal Component Analysis (PCA) of predicted KEGG Orthologs (KOs) reveals comparatively weaker differentiation between host species. **(C)** MaAsLin2 modeling of PICRUSt2-inferred KOs identified the most differentiated functional pathways between *Eulemur rufifrons* and *E. cinereiceps* relative to hybrids. Rows represent significantly associated KOs (FDR *q* < 0.05), colored by Log₂ fold-change estimates. Positive values (yellow–orange) indicate enrichment and negative values (blue–purple) indicate depletion relative to hybrids. Functions related to carbohydrate and lipid metabolism, membrane transport, stress response, and microbial defense systems exhibit strong host-specific functional differentiation.

In contrast to the clear signal of divergence among host species in terms of microbial beta diversity, predicted functional profiles exhibited broad overlap. PCA of KEGG Ortholog (KO) abundances formed one largely overlapping cluster encompassing *E. rufifrons*, *E. cinereiceps*, and hybrids (Fig. 3B). Nonetheless, we found a weak but significant effect of host species identity on KO abundance (R^2^ = 0.06; *F-value* = 3.05; *p-value* = 0.012); of the 13,239 KOs recovered in the dataset, the abundances of 338 differed significantly between hybrids and one or both parental species (threshold: adjusted *p-value* < 0.05; top 50 KOs visualized in Fig. 3C). Of these significant KOs, the vast majority (96%) reflected comparisons between *E. cinereiceps* and hybrids. Categorized broadly, most significant KOs were associated with genetic information processing (43%), metabolism (37%), cellular processes (11%), and environmental information processing (8%). KOs involved in metabolism, prokaryotic defense, and cellular interactions were particularly enriched in *E. cinereiceps* (Fig. 3C; Table S9).

## DISCUSSION

We are still in the early stages of understanding how holobionts evolve as a collective organism. Hosts and microbial symbionts face different selective pressures and evolve on vastly different timescales. Yet, their evolutionary trajectories are intertwined, particularly in mammals, which have evolved species-specific gut microbial assemblages responsible for a wide range of biological functions. Pairing microbiome data with fine-scale population histories clarifies how, when, and why hosts evolve unique microbiome communities. Here, we presented a case study using a purported hybrid species of brown lemur in southeastern Madagascar. Though our population genomic data did not support a hypothesis of hybrid speciation, some lines of evidence indicated hybrids have begun to diverge from parental species (e.g., low interclass heterozygosity). Thus, this system offers a window into the evolution of gut microbiome specificity in the face of interspecific gene flow. We found that even given shallow host divergence and substantial admixture, the gut microbiome segregated by host “species.” Below, we discuss insights into microbiome assembly, the evolutionary history of the *E. rufifrons* x *E. cinereiceps* hybrid zone, and how we can use this information to enhance conservation efforts.

### The hybrid gut microbiome is predominantly shaped by host ancestry

In this *Eulemur* system, we found that evolutionary history had a stronger influence than ecology in shaping the hybrid gut microbiome. Hybrid gut microbiomes were intermediate to parentals despite significant ecological niche divergence [53]. However, the hybrid zone is positioned between *E. rufifrons* in the north and *E. cinereiceps* in the south, raising the possibility that geography confounds evolutionary signal. To isolate the effects of each variable, we used Partial Mantel tests, ultimately finding that genomic distance had a stronger effect than both geography and environment (Table S8). This is further supported by the finding that hybrid gut microbiomes had greater compositional similarity with *E. rufifrons* than *E. cinereiceps*, reciprocating patterns of biased *E. rufifrons* ancestry in our population genomic analyses (Fig. 1A-D). Thus, the gut microbiome can reflect fine-scale population structure and gene flow.

The emergence of population-specific signatures indicates microbiomes segregate early in the speciation process. We hypothesized this divergence would be driven by the accumulation of novel microbiota in each host lineage. However, our 4H analysis showed that hybrid microbiomes were characterized by the loss, not addition, of core microbes relative to parentals (Fig. 2C). This may reflect lesser-known processes driving primate gut microbiome variation, including host control via the immune system [83] and/or microbe-microbe interactions [84, 85]. One possibility is that hybrids contain alleles from both parental immune systems, which combine to create a stronger microbial filter [43].

Despite this taxonomic depletion in hybrids, overall microbial function was largely conserved across hosts (Fig. 3B; Fig. S2), suggesting that, on a community-scale, the preservation of innate host functioning trumps ecological innovation in the hybrid gut microbiome. However, clear ecological signals were detected in the differential abundance of predicted metabolic pathways linked to diet. Relative to hybrids, *E. cinereiceps* was enriched for nutrient acquisition from structurally complex plant materials (e.g., amino acids, carbohydrates, glycan, and nucleotides). Both hybrids and *E. rufifrons* were enriched for the metabolism of simple sugars, namely fructose and mannose (Table S9). These patterns indicate a higher degree of frugivory in hybrids and *E. rufifrons* compared to *E. cinereiceps* [86, 87] — a finding which has been confirmed with behavioral observations [50, 88].

Just a few other studies have examined gut microbiome variation across natural hybrid zones. Though results have varied across systems, most studies have detected a strong role of host ancestry in shaping the gut microbiomes of hybrid mammals [36, 37, 89, 90, 91]; but see [43, 92]. In a few cases, hybrid gut microbiota more closely resembled either the matrilineal [37, 91] or paternal species [90]. In at least one example, the strong effects of ancestry created a diet-microbiome mismatch; despite reliance on specialized gut bacteria for the detoxification of plant secondary compounds in certain habitats, hybrid microbiomes were shaped by host genotype, not habitat or diet [36]. Taken together, these insights into the hybrid microbiome suggest that in many hosts, the limits of gut microbiome divergence are bounded by phylogenetic inertia. However, despite being intermediate, hybrid gut microbiomes were clearly distinct from parentals in our study and others, indicating that phylosymbiosis can arise early in the speciation process, even when species boundaries are leaky.

One caveat to our study is that, because low-coverage host genome sequencing required as much source material as possible, microbiome and genomic data were not always derived from the same samples. We mitigated this incongruity by selecting demographically similar individuals from the same social group, but our results should not be interpreted as one-to-one correlations.

### A revised evolutionary history of E. rufifrons x E. cinereiceps hybridization

Our results confirm a history of hybridization between *E. rufifrons* and *E. cinereiceps*. We also confirm reports that the hybrid zone appears to be old and stable [52,53], as evidenced by the absence of F1 and F2 hybrids and low interclass heterozygosity of all hybrid individuals (Fig. 1D). This suggests an advanced-generation hybrid zone and/or that hybrids have been mating with other hybrids. Nonetheless, our data do not support a hypothesis of hybrid speciation. Rather than forming a discrete third genetic cluster, hybrid individuals exhibited a continuum of ancestry proportions between the two parental species (Fig. 1A–C). The absence of F1 and F2 hybrids, coupled with widespread advanced-generation admixture, suggests that hybridization has persisted over many generations but has not resulted in the emergence of a reproductively isolated hybrid lineage. Instead, the genetic data are more consistent with a long-standing hybrid zone.

The two population clusters correspond with parental species identifications, with *E. cinereiceps* in the south, *E. rufifrons* in the north, and extensive admixture in the middle. Nearly all individuals sampled in the hybrid zone contained genetic material from both parental species, but contributions from *E. rufifrons* exceeded *E. cinereiceps* (Fig. 1A–D; Table S3). Though the hybrid zone was largely restricted to eastern Andrinigtra National Park, hybrids were detected as far north as Ranomafana National Park. Admixture south of the hybrid zone was less pervasive, suggesting a history of biased introgression and relative distinctiveness of *E. cinereiceps*. The Emma site represented a boundary between *E. cinereiceps* and the hybrid zone (Fig. 1A–D). Corresponding botanical analyses indicated Emma’s forest structure was highly dissimilar compared to other sites sampled in this study (unpublished data), raising the possibility that environmental selection may have restricted the range of either *E. cinereiceps* or hybrids. Lending additional support for this idea, we detected a significant signal of IBE between *E. cinereiceps* and hybrids (Table S4, S5). This contrasts with our finding of IBD, not IBE, between *E. rufifrons* and hybrids (Table S4, S5). Thus, the northern extent of the hybrid zone appears to reflect dispersal limitation, not environmental selection.

### Conservation implications

This study examined hybridization between two primate species threatened with extinction. The IUCN Red List (2020) classifies *E. rufifrons* as Vulnerable and *E. cinereiceps* as Critically Endangered. Hybridization has complicated efforts to accurately delineate the geographic distributions of these taxa. By identifying the limits of the hybrid zone, our findings provide a clearer framework for studying and conserving these species.

Our results indicate that *E. cinereiceps* is largely restricted to the southern COFAV, occupying a narrower and lower-elevation range than either *E. rufifrons* or hybrids. These environmental characteristics contribute to its heightened endangerment. Low elevation forests are particularly susceptible to conversion for agriculture [93, 94, 95], and the narrow distribution of suitable habitat increases the likelihood of human-lemur encounters.

These threats were apparent during field sampling. We found and dismantled 26 bushmeat traps targeting lemurs, all of which were located in Emma (unpublished data). Three of these traps contained live lemurs from the same social group: an adult female, an adult male, and a juvenile female. Genetic analyses performed in this study confirmed that the juvenile female belonged to *E. cinereiceps*, underscoring the direct and ongoing hunting pressure this species faces. As a next step, we recommend targeted conservation intervention in the southern COFAV, particularly Emma, to ensure the survival of *E. cinereiceps*. The most effective conservation strategies in southeastern Madagascar have integrated conservation education, reforestation, sustainable agriculture, and access to rural healthcare [96].

## DATA AVAILABILITY STATEMENT

Raw sequences are available under NCBI Bioproject PRJNA1336526. All scripts are publicly available at https://github.com/katherinermartin/Donohue_et_al.

## CONFLICT OF INTEREST

Patricia C. Wright is on the advisory board of Primate Conservation Inc, one of the funders of this project. She did not advise on funding this project.

## ACKNOWLEDGEMENTS

We thank the Ministry of Environment and Sustainable Development for permitting this research, RAKOTOZAFY Joseph and RAVELOMANAZAFY Le vaillant Arsène for sharing their local knowledge, Centre ValBio and MICET for supporting our field expeditions, and Dr. Steig Johnson for his advice and foundational information about the hybrid system. We acknowledge funding support from NSF #2207198 (DWW and KME), Fulbright U.S. Student Program (MED), Primate Conservation Inc. (MED), University of Kentucky’s Dean’s Competitive Fellowship (MED), University of Kentucky’s Morgan Fellowship (MED), and University of Kentucky’s Ribble Mini Research Grant (MED).

## Notes

https://github.com/katherinermartin/Donohue_et_al

### Summary of Updates

Expanded gut microbiome analyses with significant revisions to the main text.

## WORKS CITED

[1] Bordenstein, S. R., & Theis, K. R. (2015). Host biology in light of the microbiome: Ten principles of Holobionts and hologenomes. PLOS Biology, 13(8), e1002226. 10.1371/journal.pbio.1002226

[2] Hooper, L. V., Littman, D. R., & Macpherson, A. J. (2012). Interactions between the microbiota and the immune system. Science, 336(6086), 1268–1273. 10.1126/science.1223490

[3] Hacquard, S., Garrido-Oter, R., González, A., Spaepen, S., Ackermann, G., Lebeis, S., McHardy, A. C., Dangl, J. L., Knight, R., Ley, R., & Schulze-Lefert, P. (2015). Microbiota and host nutrition across plant and animal kingdoms. Cell Host & Microbe, 17(5), 603–616. 10.1016/j.chom.2015.04.009

[4] Voolstra, C. R., & Ziegler, M. (2020). Adapting with microbial help: Microbiome flexibility facilitates rapid responses to environmental change. BioEssays, 42(7), 2000004. 10.1002/bies.202000004

[5] Margulis, L., & René Fester. (1991). Symbiosis as a source of evolutionary innovation: Speciation an morphogenesis. Mit Press.

[6] Amato, K. R., Yeoman, C. J., Kent, A., Righini, N., Carbonero, F., Estrada, A., Rex Gaskins, H., Stumpf, R. M., Yildirim, S., Torralba, M., Gillis, M., Wilson, B. A., Nelson, K. E., White, B. A., & Leigh, S. R. (2013). Habitat degradation impacts black howler monkey (*Alouatta pigra*) gastrointestinal microbiomes. The ISME Journal, 7(7), 1344–1353. 10.1038/ismej.2013.16

[7] Barelli, C., Albanese, D., Donati, C., Pindo, M., Dallago, C., Rovero, F., Cavalieri, D., Tuohy, K. M., Hauffe, H. C., & De Filippo, C. (2015). Habitat fragmentation is associated to gut microbiota diversity of an endangered primate: Implications for conservation. Scientific Reports, 5(1). 10.1038/srep14862

[8] Wu, Y., Yang, Y., Cao, L., Yin, H., Xu, M., Wang, Z., Liu, Y., Wang, X., & Deng, Y. (2018). Habitat environments impacted the gut microbiome of long-distance migratory swan geese but central species conserved. Scientific Reports, 8(1). 10.1038/s41598-018-31731-9

[9] Shankregowda, A. M., Siriyappagouder, P., Kuizenga, M., Bal, T. M. P., Abdelhafiz, Y., Eizaguirre, C., Fernandes, J. M. O., Kiron, V., & Raeymaekers, J. A. M. (2023). Host habitat rather than evolutionary history explains gut microbiome diversity in sympatric stickleback species. Frontiers in Microbiology, 14. 10.3389/fmicb.2023.1232358

[10] Wu, G. D., Chen, J., Hoffmann, C., Bittinger, K., Chen, Y.-Y., Keilbaugh, S. A., Bewtra, M., Knights, D., Walters, W. A., Knight, R., Sinha, R., Gilroy, E., Gupta, K., Baldassano, R., Nessel, L., Li, H., Bushman, F. D., & Lewis, J. D. (2011). Linking long-term dietary patterns with gut microbial enterotypes. Science, 334(6052), 105–108. 10.1126/science.1208344

[11] Muegge, B. D., Kuczynski, J., Knights, D., Clemente, J. C., Gonzalez, A., Fontana, L., Henrissat, B., Knight, R., & Gordon, J. I. (2011). Diet drives convergence in gut microbiome functions across mammalian phylogeny and within humans. Science, 332(6032), 970–974. 10.1126/science.1198719

[12] Turnbaugh, P. J., Ridaura, V. K., Faith, J. J., Rey, F. E., Knight, R., & Gordon, J. I. (2009). The effect of diet on the human gut microbiome: A metagenomic analysis in humanized gnotobiotic mice. Science Translational Medicine, 1(6), 6ra14–6ra14. 10.1126/scitranslmed.3000322

[13] David, L. A., Maurice, C. F., Carmody, R. N., Gootenberg, D. B., Button, J. E., Wolfe, B. E., Ling, A. V., Devlin, A. S., Varma, Y., Fischbach, M. A., Biddinger, S. B., Dutton, R. J., & Turnbaugh, P. J. (2013). Diet rapidly and reproducibly alters the human gut microbiome. Nature, 505(7484), 559–563. 10.1038/nature12820

[14] Brooks, A. W., Kohl, K. D., Brucker, R. M., van Opstal, E. J., & Bordenstein, S. R. (2016). Phylosymbiosis: Relationships and functional effects of microbial communities across host evolutionary history. PLOS Biology, 14(11), e2000225. 10.1371/journal.pbio.2000225

[15] Beasley, D. E., Koltz, A. M., Lambert, J. E., Fierer, N., & Dunn, R. R. (2015). The evolution of stomach acidity and its relevance to the human microbiome. PLOS One, 10(7). 10.1371/journal.pone.0134116

[16] Douglas, A.E. & Werren, J.H. 2016. Holes in the Hologenome: Why Host-Microbe Symbioses Are Not Holobionts. mBio, 7:10.1128/mbio.02099-15. 10.1128/mbio.02099-15

[17] Mazel, F., Davis, K. M., Loudon, A., Kwong, W. K., Groussin, M., & Parfrey, L. W. (2018). Is host filtering the main driver of phylosymbiosis across the tree of Life? mSystems, 3(5). 10.1128/msystems.00097-18

[18] Amato, K. R., Mallott, E. K., McDonald, D., Dominy, N. J., Goldberg, T., Lambert, J. E., Swedell, L., Metcalf, J. L., Gomez, A., Britton, G. A. O., Stumpf, R. M., Leigh, S. R., & Knight, R. (2019). Convergence of human and old world monkey gut microbiomes demonstrates the importance of human ecology over phylogeny. Genome Biology, 20(1). 10.1186/s13059-019-1807-z

[19] Moeller, A. H., Suzuki, T. A., Lin, D., Lacey, E. A., Wasser, S. K., & Nachman, M. W. (2017). Dispersal limitation promotes the diversification of the mammalian gut microbiota. Proceedings of the National Academy of Sciences, 114(52), 13768–13773. 10.1073/pnas.1700122114

[20] Mazel, F., Guisan, A., & Parfrey, L. W. (2023). Transmission mode and dispersal traits correlate with host specificity in mammalian gut microbes. Molecular Ecology, 33(1). 10.1111/mec.16862

[21] Gaulke, C. A., Arnold, H. K., Humphreys, I. R., Kembel, S. W., O’Dwyer, J. P., & Sharpton, T. J. (2018). Ecophylogenetics clarifies the evolutionary association between mammals and their gut microbiota. mBio, 9(5). 10.1128/mbio.01348-18

[22] Groussin, M., Mazel, F., Sanders, J. G., Smillie, C. S., Lavergne, S., Thuiller, W., & Alm, E. J. (2017). Unraveling the processes shaping mammalian gut microbiomes over evolutionary time. Nature Communications, 8(1). 10.1038/ncomms14319

[23] Koskella, B., & Bergelson, J. (2020). The study of host–microbiome (co)evolution across levels of selection. Philosophical Transactions of the Royal Society B: Biological Sciences, 375(1808), 20190604. 10.1098/rstb.2019.0604

[24] Mallott, E. K., & Amato, K. R. (2021). Host specificity of the gut microbiome. Nature Reviews Microbiology, 19(10), 639–653. 10.1038/s41579-021-00562-3

[25] Coyne, H. A., & H Allen Orr. (2004). Speciation. Sinauer Associates, Cop.

[26] Smadja, C.M., & Butlin, R.K. (2011). A framework for comparing processes of speciation in the presence of gene flow. Molecular Ecology, 20(24), 5123–5140. 10.1111/j.1365-294x.2011.05350.x

[27] Feder, J. L., Egan, S. P., & Nosil, P. (2012). The genomics of speciation-with-gene-flow. Trends in Genetics, 28(7), 342–350. 10.1016/j.tig.2012.03.009

[28] Martin, S. H., Dasmahapatra, K. K., Nadeau, N. J., Salazar, C., Walters, J. R., Simpson, F., Blaxter, M., Manica, A., Mallet, J., & Jiggins, C. D. (2013). Genome-wide evidence for speciation with gene flow in *Heliconius* butterflies. Genome Research, 23(11), 1817–1828. 10.1101/gr.159426.113

[29] Fraïsse, C., Roux, C., Welch, J. J., & Bierne, N. (2014). Gene-Flow in a mosaic hybrid zone: Is local introgression adaptive? Genetics, 197(3), 939–951. 10.1534/genetics.114.161380

[30] Tigano, A., & Friesen, V. L. (2016). Genomics of local adaptation with gene flow. Molecular Ecology, 25(10), 2144–2164. 10.1111/mec.13606

[31] Reilly, P. F., Tjahjadi, A., Miller, S. L., Akey, J. M., & Tucci, S. (2022). The contribution of neanderthal introgression to modern human traits. Current Biology, 32(18), R970–R983. 10.1016/j.cub.2022.08.027

[32] Mallet, J. (2007). Hybrid speciation. Nature, 446(7133), 279–283. 10.1038/nature05706

[33] Abbott, R. J., & Rieseberg, L. H. (2021). Hybrid speciation. Encyclopedia of Life Sciences, 1–9. 10.1002/9780470015902.a0029379

[34] Orr, H. A., & Turelli, M. (2001). The evolution of postzygotic isolation: Accumulating Dobzhansky-Muller Incompatibilities. Evolution, 55(6), 1085–1094. 10.1111/j.0014-3820.2001.tb00628.x

[35] Price, T. D., & Bouvier, M.M. (2002). The evolution of F_1_postzygotic incompatibilities in birds. Evolution, 56(10), 2083–2089. 10.1111/j.0014-3820.2002.tb00133.x

[36] Nielsen, D. P., Harrison, J. G., Byer, N. W., Faske, T. M., Parchman, T. L., Simison, W. B., & Matocq, M. D. (2022). The gut microbiome reflects ancestry despite dietary shifts across a hybrid zone. Ecology Letters, 26(1), 63–75. 10.1111/ele.14135

[37] Abraham, J. O., Lin, B., Miller, A. E., Henry, L. P., Demmel, M. Y., Warungu, R., Mwangi, M., Lobura, P. M., Pallares, L. F., Ayroles, J. F., Pringle, R. M., & Rubenstein, D. I. (2024). Determinants of microbiome composition: Insights from free-ranging hybrid zebras (*Equus quagga × grevyi*). Molecular Ecology, 33(11). 10.1111/mec.17370

[38] Russell, A. C., Vohsen, S. A., Spinelli, J. M. C., Martinez, N., Curry, R. L., Roth, T. C., Semenov, G. A., Taylor, S. A., & Rice, A. M. (2026). Environment impacts gut microbiome richness in hybridizing chickadees, while ancestry influences community composition. Ecosphere, 17(4). 10.1002/ecs2.70611

[39] Camper, B. T., Laughlin, Z., Malagon, D., Denton, R., & Bewick, S. (2024). A conceptual framework for host-associated microbiomes of hybrid organisms. Methods in Ecology and Evolution, 15(3), 511–529. 10.1111/2041-210x.14279

[40] Camper, B. T., Kanes, A. S., Laughlin, Z. T., Manuel, R. T., & Bewick, S. A. (2025). Transgressive hybrids as hopeful holobionts. Microbiome, 13(1). 10.1186/s40168-024-01994-8

[41] Theißen, G. (2009). Saltational evolution: Hopeful monsters are here to stay. Theory in Biosciences, 128(1), 43–51. 10.1007/s12064-009-0058-z

[42] Brucker, R. M., & Bordenstein, S. R. (2012). Speciation by symbiosis. Trends in Ecology & Evolution, 27(8), 443–451. 10.1016/j.tree.2012.03.011

[43] Wang, J., Kalyan, S., Steck, N., Turner, L. M., Harr, B., Künzel, S., Vallier, M., Häsler, R., Franke, A., Oberg, H.-H., Ibrahim, S. M., Grassl, G. A., Kabelitz, D., & Baines, J. F. (2015). Analysis of intestinal microbiota in hybrid house mice reveals evolutionary divergence in a vertebrate hologenome. Nature Communications, 6(1). 10.1038/ncomms7440

[44] Wogan, G. O. U., Yuan, M. L., Luke, M. D, & Wang, I. J. (2023). Hybridization and transgressive evolution generate diversity in an adaptive radiation of Anolis lizards. Systematic Biology, 72(4), 874–884.

[45] Combrink, L. L., Jimena Golcher-Benavides, Lewanski, A. L., Rick, J. A., Rosenthal, W. C., & Wagner, C. E. (2024). Population genomics of adaptive radiation. Molecular Ecology, 34(2). 10.1111/mec.17574

[46] Everson, K. M., Pozzi, L., Barrett, M. A., Blair, M. E., Donohue, M. E., Kappeler, P. M., Kitchener, A. C., Lemmon, A. R., Lemmon, E. M., Pavón-Vázquez, C. J., Radespiel, U., Blanchard Randrianambinina, Rasoloarison, R. M., Solofonirina Rasoloharijaona, Roos, C., Jordi Salmona, Yoder, A. D, Zenil-Ferguson, R., Zinner, D., & Weisrock, D. W. (2025). Multiple bursts of speciation in Madagascar’s endangered lemurs. Nature Communications, 16(1). 10.1038/s41467-025-62310-y

[47] Orkin, J. D, Lukas, Núria Hermosilla-Albala, Fontsere, C., Aylward, M. L., Janiak, M. C., Andriaholinirina, N., Balaresque, P., Blair, M. E., Jean-Luc Fausser, Gut, I. G., Gut, M., Hahn, M. W., Harris, R. A., Horvath, J. E., Keyser, C., Kitchener, A. C., Le, M. D, Lizano, E., … Bonet, T. M. (2024). Ecological and anthropogenic effects on the genomic diversity of lemurs in Madagascar. Nature Ecology & Evolution. 10.1038/s41559-024-02596-1

[48] Everson, K. M., Donohue, M. E., & Weisrock, D. W. (2023). A pervasive history of gene flow in Madagascar’s true lemurs (genus *Eulemur*). Genes, 14(6), 1130–1130. 10.3390/genes14061130

[49] Mercuri, G., Merici, G., Farh, K. K.-H., Lukas F.K. Kuderna, Rogers, J., Tomàs Marques- Bonet, Donati, G., Riccardo Percudani, & Capelli, C. (2025). Leaping between branches: Hybridisation and the tangled evolutionary history of true lemurs. Molecular Phylogenetics and Evolution, 215, 108503–108503. 10.1016/j.ympev.2025.108503

[50] Johnson, S. E. (2002). Ecology and speciation in Brown lemurs, white-collared lemurs (Eulemur albocollaris) and hybrids (Eulemur albocollaris x Eulemur fulvus rufus) in southeastern madagascar. The University of Texas at Austin.

[51] Wyner, Y. M., Johnson, S. E., Stumpf, R. M., & Desalle, R. (2002). Genetic assessment of a white-collared×red-fronted lemur hybrid zone at Andringitra, Madagascar. American Journal of Primatology, 57(2), 51–66. 10.1002/ajp.10033

[52] Delmore, K. E., Brenneman, R. A., Lei, R., Bailey, C. A., Brelsford, A., Louis, E. E., & Johnson, S. E. (2013). Clinal variation in a brown lemur (*Eulemur* spp.) hybrid zone: Combining morphological, genetic and climatic data to examine stability. Journal of Evolutionary Biology, 26(8), 1677–1690. 10.1111/jeb.12178

[53] Johnson, S. E., Delmore, K. E., Brown, K. A., Wyman, T. M., & Louis, E. E. (2015). Niche divergence in a brown lemur (*Eulemur* spp.) Hybrid zone: Using ecological niche models to test models of Stability. International Journal of Primatology, 37(1), 69–88. 10.1007/s10764-015-9872-y

[54] Batist, C. H., Bliss, M., Weisenbeck, D. R., Rafamantanantsoa, R., Razafindrakoto, G., Ranaivosolo, J. W., Baptiste, V. J., Wright, P. C., & Donohue, M. E. (2023). Updated lemur species ranges in Madagascar’s Corridor Forestier d’Ambositra Vondrozo (COFAV). Folia Primatologica, 94(4–6), 207–223. 10.1163/14219980-bja10012

[55] Chiou, K. L., & Bergey, C. M. (2018). Methylation-based enrichment facilitates low-cost, noninvasive genomic scale sequencing of populations from feces. Scientific Reports, 8(1). 10.1038/s41598-018-20427-9

[56] Kozich, J. J., Westcott, S. L., Baxter, N. T., Highlander, S. K., & Schloss, P. D. (2013). Development of a dual-index sequencing strategy and curation pipeline for analyzing amplicon sequence data on the MiSeq illumina sequencing platform. Applied and Environmental Microbiology, 79(17), 5112–5120. 10.1128/aem.01043-13

[57] Korneliussen, T. S., Albrechtsen, A., & Nielsen, R. (2014). angsd: analysis of next generation sequencing data. BMC Bioinformatics, 15(1). 10.1186/s12859-014-0356-4

[58] Meisner, J., & Albrechtsen, A. (2018). Inferring population structure and admixture proportions in low-depth NGS data. Genetics, 210(2), 719–731. 10.1534/genetics.118.301336

[59] Wickham, H. (2016). Ggplot2: Elegant graphics for data analysis. In Use R! Springer International Publishing. 10.1007/978-3-319-24277-4

[60] Skotte, L., Korneliussen, T. S., & Albrechtsen, A. (2013). Estimating individual admixture proportions from Next generation sequencing data. Genetics, 195(3), 693–702. 10.1534/genetics.113.154138

[61] Kopelman, N. M., Mayzel, J., Jakobsson, M., Rosenberg, N. A., & Mayrose, I. (2015). Clumpak : A program for identifying clustering modes and packaging population structure inferences across k. Molecular Ecology Resources, 15(5), 1179–1191. 10.1111/1755-0998.12387

[62] Evanno, G., Regnaut, S., & Goudet, J. (2005). Detecting the number of clusters of individuals using the software structure: A simulation study. Molecular Ecology, 14(8), 2611– 2620. 10.1111/j.1365-294x.2005.02553.x

[63] Jakobsson, M., & Rosenberg, N. A. (2007). Clumpp: A cluster matching and permutation program for dealing with label switching and multimodality in analysis of population structure. Bioinformatics, 23(14), 1801–1806. 10.1093/bioinformatics/btm233

[64] Francis, R. M. (2016). pophelper: An r package and web app to analyse and visualize population structure. Molecular Ecology Resources, 17(1), 27–32. 10.1111/1755-0998.12509

[65] Wiens, B. J., DeCicco, L. H., & Colella, J. P. (2025). triangulaR: An r package for identifying AIMs and building triangle plots using SNP data from hybrid zones. Heredity, 134(5), 251–262. 10.1038/s41437-025-00760-2

[66] Wiens, B. J., & Colella, J. P. (2024). That is not a hybrid: How to distinguish patterns of admixture and isolation by distance. Molecular Ecology Resources, 25(3). 10.1111/1755-0998.14039

[67] Wang, I. J. (2013). Examining the full effects of landscape heterogeneity on spatial genetic variation: A multiple matrix regression approach for quantifying geographic and ecological isolation. Evolution, 67(12), 3403–3411. 10.1111/evo.12134

[68] Ferrier, S., Manion, G., Elith, J., & Richardson, K. (2007). Using generalized dissimilarity modelling to analyse and predict patterns of beta diversity in regional biodiversity assessment. Diversity and Distributions, 13(3), 252–264. 10.1111/j.1472-4642.2007.00341.x

[69] Fick, S. E., & Hijmans, R. J. (2017). WorldClim 2: New 1-km spatial resolution climate surfaces for global land areas. International Journal of Climatology, 37(12), 4302–4315.

[70] Chambers, E.A., Bishop, A. P., & Wang, I. J. (2023). Individual-based landscape genomics for conservation: An analysis pipeline. Molecular Ecology Resources. 10.1111/1755-0998.13884

[71] Bolyen, E., Rideout, J. R., Dillon, M. R., Bokulich, N. A., Abnet, C. C., Al-Ghalith, G. A., Alexander, H., Alm, E. J., Arumugam, M., Asnicar, F., Bai, Y., Bisanz, J. E., Bittinger, K., Brejnrod, A., Brislawn, C. J., Brown, C. T., Callahan, B. J., Caraballo-Rodríguez, A. M., Chase, J., … Caporaso, J. G. (2019). Reproducible, interactive, scalable and extensible microbiome data science using qiime 2. Nature Biotechnology, 37(8), 852–857. 10.1038/s41587-019-0209-9

[72] Callahan, B. J., McMurdie, P. J., Rosen, M. J., Han, A. W., Johnson, A. J. A., & Holmes, S. P. (2016). DADA2: High-resolution sample inference from Illumina amplicon data. Nature Methods, 13(7), 581–583. 10.1038/nmeth.3869

[73] Kruskal, W. H., & Wallis, W. A. (1952). Use of ranks in one-criterion variance analysis. Journal of the American Statistical Association, 47(260), 583–621. 10.1080/01621459.1952.10483441

[74] Conover, W. J., Johnson, M. E., & Johnson, M. M. (1981). A comparative study of tests for Homogeneity of Variances, with applications to the outer continental shelf bidding data. Technometrics, 23(4), 351–361. 10.1080/00401706.1981.10487680

[75] Kanehisa, M., Sato, Y., Kawashima, M., Furumichi, M., & Tanabe, M. (2015). KEGG as a reference resource for gene and protein annotation. Nucleic Acids Research, 44(D1), D457–D462. 10.1093/nar/gkv1070

[76] Douglas, G. M., Maffei, V. J., Zaneveld, J. R., Yurgel, S. N., Brown, J. R., Taylor, C. M., Huttenhower, C., & Langille, M. G. I. (2020). PICRUSt2 for prediction of metagenome functions. Nature Biotechnology, 38(6), 685–688. 10.1038/s41587-020-0548-6

[77] Yang, C., Mai, J., Cao, X., Burberry, A., Fabio Cominelli, & Zhang, L. (2023). Ggpicrust2: An r package for PICRUSt2 predicted functional profile analysis and visualization. Bioinformatics, 39(8). 10.1093/bioinformatics/btad470

[78] Dixon, P. (2003). Vegan, a package of r functions for community ecology. Journal of Vegetation Science, 14(6), 927–930. 10.1111/j.1654-1103.2003.tb02228.x

[79] Mallick, H., Rahnavard, A., McIver, L. J., Ma, S., Zhang, Y., Nguyen, L. H., Tickle, T. L., Weingart, G., Ren, B., Schwager, E. H., Chatterjee, S., Thompson, K. N., Wilkinson, J. E., Subramanian, A., Lu, Y., Waldron, L., Paulson, J. N., Franzosa, E. A., Bravo, H. C., & Huttenhower, C. (2021). Multivariable association discovery in population-scale meta-omics studies. PLOS Computational Biology, 17(11), e1009442. 10.1371/journal.pcbi.1009442

[80] Kanehisa, M., & Goto, S. (2000). KEGG: Kyoto encyclopedia of genes and genomes. Nucleic Acids Research, 28(1), 27–30.

[81] Gomez, A., Petrzelkova, K., Yeoman, C. J., Vlckova, K., Mrázek, J., Koppova, I., Carbonero, F., Ulanov, A., Modry, D., Todd, A., Torralba, M., Nelson, K. E., Gaskins, H. R., Wilson, B., Stumpf, R. M., White, B. A., & Leigh, S. R. (2015). Gut microbiome composition and metabolomic profiles of wild western lowland gorillas (*Gorilla gorilla gorilla*) reflect host ecology. Molecular Ecology, 24(10), 2551–2565. 10.1111/mec.13181

[82] Clayton, J. B., Al-Ghalith, G. A., Long, H. T., Tuan, B. Van, Cabana, F., Huang, H., Vangay, P., Ward, T., Minh, V. Van, Tam, N. A., Dat, N. T., Travis, D. A., Murtaugh, M. P., Covert, H., Glander, K. E., Nadler, T., Toddes, B., Sha, J. C. M., Singer, R., … Johnson, T. J. (2018). Associations between nutrition, gut microbiome, and health in a novel nonhuman primate model. Scientific Reports, 8(1). 10.1038/s41598-018-29277-x

[83] Wilde, J., Slack, E., & Foster, K. R. (2024). Host control of the microbiome: Mechanisms, evolution, and disease. Science, 385(6706). 10.1126/science.adi3338

[84] Coyte, K. Z., & Rakoff-Nahoum, S. (2019). Understanding competition and cooperation within the mammalian gut microbiome. Current Biology, 29(11), R538–R544. 10.1016/j.cub.2019.04.017

[85] Palmer, J. D., & Foster, K. R. (2022). Bacterial species rarely work together. Science, 376(6593), 581–582. 10.1126/science.abn5093

[86] Amato, K. R., Sanders, J. G., Song, S. J., Nute, M., Metcalf, J. L., Thompson, L. R., Morton, J. T., Amir, A., McKenzie, V. J., Humphrey, G., Gogul, G., Gaffney, J., Baden, A. L., Britton, G. A. O., Cuozzo, F. P., Di Fiore, A., Dominy, N. J., Goldberg, T. L., Gomez, A., … Leigh, S. R. (2018). Evolutionary trends in host physiology outweigh dietary niche in structuring primate gut microbiomes. The ISME Journal, 13(3), 576–587. 10.1038/s41396-018-0175-0

[87] Greene, L. K., Williams, C. V., Junge, R. E., Mahefarisoa, K. L., Rajaonarivelo, T., Rakotondrainibe, H., O’Connell, T. M., & Drea, C. M. (2020). A role for gut microbiota in host niche differentiation. The ISME Journal, 14(7), 1675–1687. 10.1038/s41396-020-0640-4

[88] Overdorff, D. J. (1993). Similarities, differences, and seasonal patterns in the diets of *Eulemur rubriventer* and *Eulemur fulvus rufus* in the Ranomafana National Park, Madagascar. International Journal of Primatology, 14(5), 721–753. 10.1007/bf02192188

[89] Malukiewicz, J., Cartwright, R. A., Dergam, J. A., Igayara, C. S., Kessler, S. E., Moreira, S. B., Nash, L. T., Nicola, P. A., Pereira, L. C. M., Pissinatti, A., Ruiz-Miranda, C. R., Ozga, A. T., Quirino, A. A., Roos, C., Silva, D. L., Stone, A. C., & Grativol, A. D. (2022). The gut microbiome of exudivorous marmosets in the wild and captivity. Scientific Reports, 12(1). 10.1038/s41598-022-08797-7

[90] Wei, L., Zeng, B., Li, B., Guo, W., Mu, Z., Gan, Y., & Li, Y. (2024). Hybridization alters red deer gut microbiome and metabolites. Frontiers in Microbiology, 15. 10.3389/fmicb.2024.1387957

[91] Zubillaga-Martín, D., Solórzano-García, B., Yanez-Montalvo, A., de León-Lorenzana, A., Falcón, L. I., & Vázquez-Domínguez, E. (2025). Gut microbiota signatures of the three mexican primate species, including hybrid populations. PLOS One, 20(3), e0317657. 10.1371/journal.pone.0317657

[92] Grieneisen, L. E., Charpentier, M. J. E., Alberts, S. C., Blekhman, R., Bradburd, G., Tung, J., & Archie, E. A. (2019). Genes, geology and germs: Gut microbiota across a primate hybrid zone are explained by site soil properties, not host species. Proceedings of the Royal Society B: Biological Sciences, 286(1901), 20190431. 10.1098/rspb.2019.0431

[93] Goodman, S. M., & Rasolonandrasana, B. P. N. (2001). Elevational zonation of birds, insectivores, rodents and primates on the slopes of the Andringitra Massif, Madagascar. Journal of Natural History, 35(2), 285–305. 10.1080/00222930150215387

[94] Deppe, A., Randriamiarisoa, M., Schütte, K., & Wright, P. (2007). A brief lemur survey of the Ranomafana Andringitra corridor region in Tolongoina, southeast Madagascar. Lemur News, 12(12).

[95] Ramiadantsoa, T., Ovaskainen, O., Rybicki, J., & Hanski, I. (2015). Large-Scale habitat corridors for biodiversity conservation: A forest corridor in Madagascar. PLOS One, 10(7), e0132126. 10.1371/journal.pone.0132126

[96] Razafindravony, L. E., Donohue, M. E., Docherty, M. A., Maggy, A. M., Lazasoa, R. S., Rafanomezantsoa, O. J. S., Ramarjaona, R. A., Randriarimanana, J. N. M., Rafanambinantsoa, A. O., Randrianarivelo, H., & Wright, P. C. (2023). Evaluating the impact of environmental education around Ranomafana National Park. American Journal of Primatology, 85(5). 10.1002/ajp.23477

