## Supplementary material for "Gene flow erodes core gut microbiota, but not microbiome function, across a natural primate hybrid zone": Fig. S1

**SUPPLEMENTAL FIGURES AND TABLES**

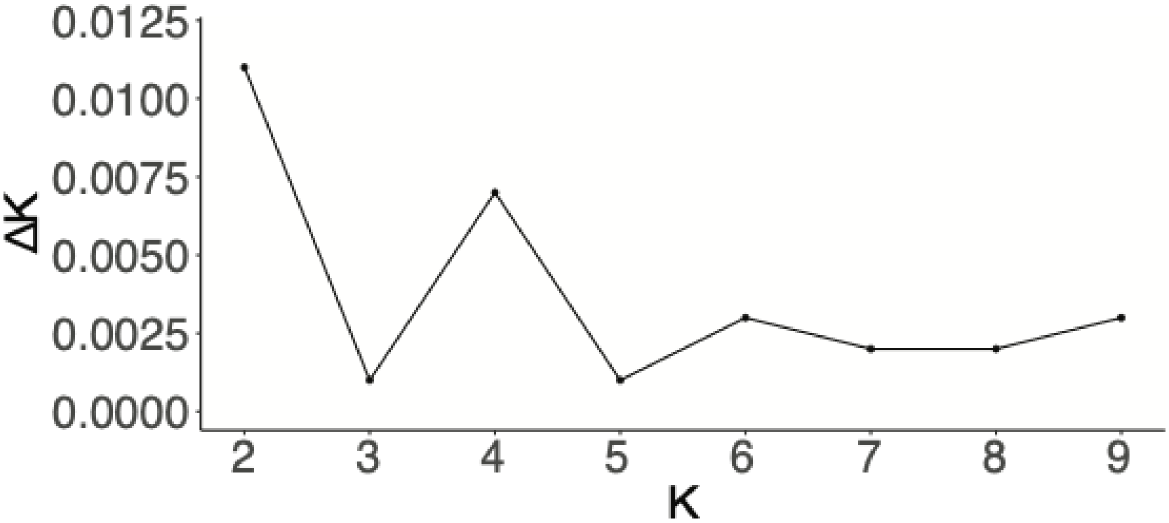


**Fig. S1:** Delta *K* distribution graph of *K* 2 through 9 generated using the Evanno method on the clumpak webserver. K *1* through *10* were evaluated as part of ngsadmix to infer the number of genetic clusters in the dataset, and the admixture proportions of each sample. Since the inferred *K* is calculated as the difference in log likelihood scores between each *K* evaluated, only *K* = 2 through *K* = *9* are shown.

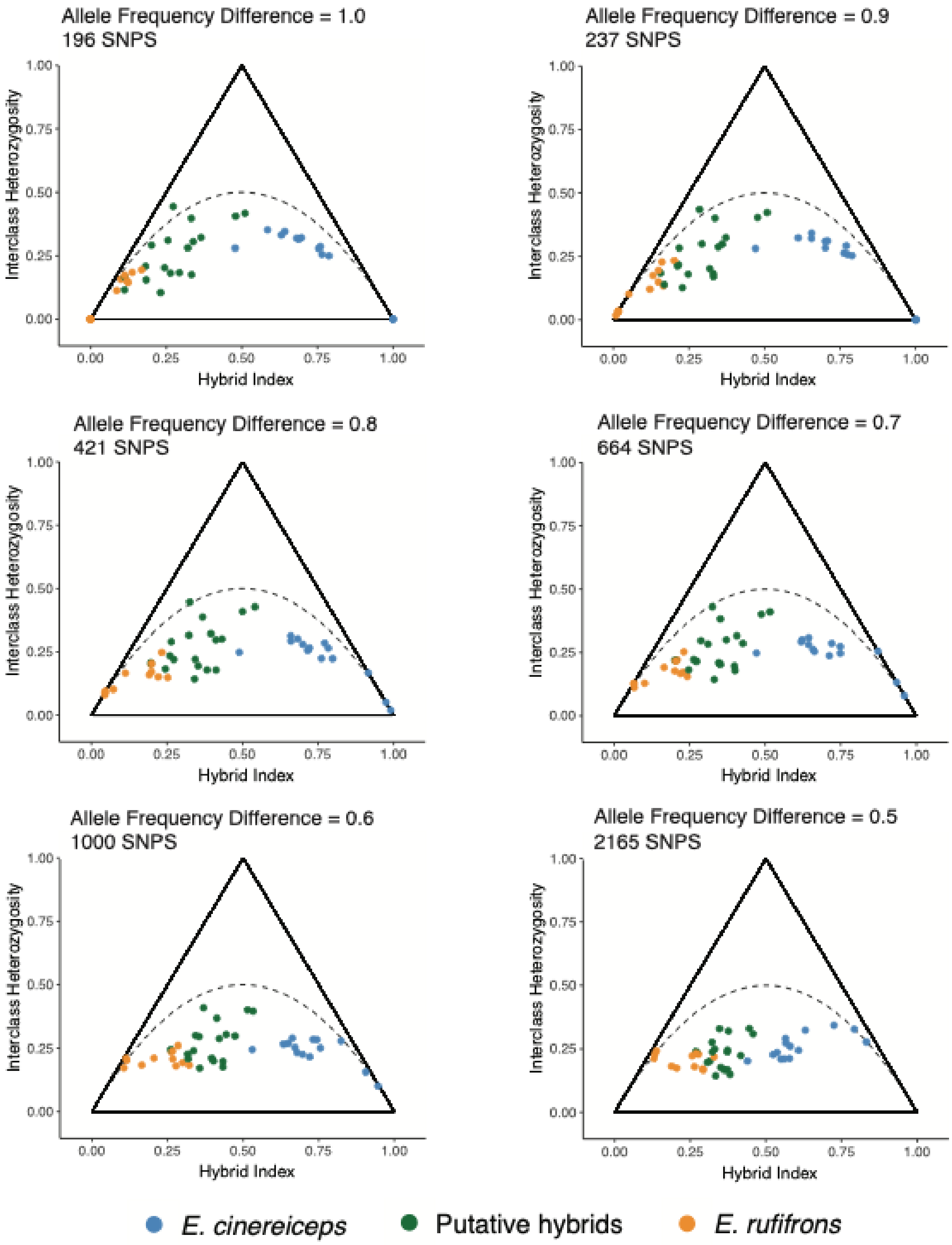


**Fig. S2:** Triangle plots showing the hybrid index (x-axis) and interclass heterozygosity (y-axis) of every individual in the dataset. Each triangle plot explored a different allele frequency difference threshold from 1.0 (top left) to 0.5 (bottom right). Note that higher allele frequency difference thresholds result in fewer total recovered SNPs. The 1.0 allele frequency difference threshold plot is also shown in the main text (Fig. 1d).

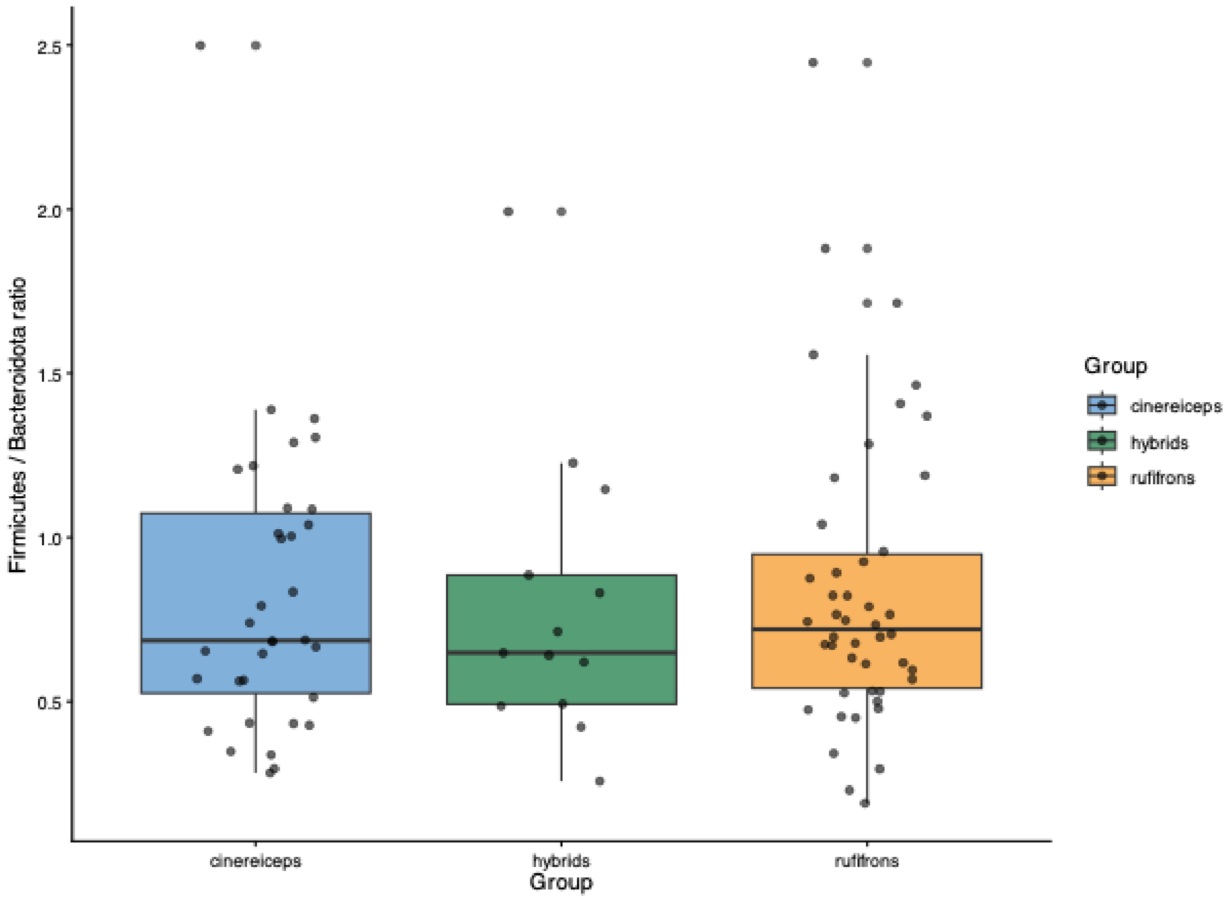


**Fig. S3:** Boxplot comparing the Firmicutes-to-Bacteroidota (F/B) ratio among *Eulemur cinereiceps*, hybrids, and *Eulemur rufifrons*. The F/B ratio did not differ significantly among the three groups (Kruskal–Wallis test: $\chi^2 = 0.204$, df = 2, $P = 0.903$).

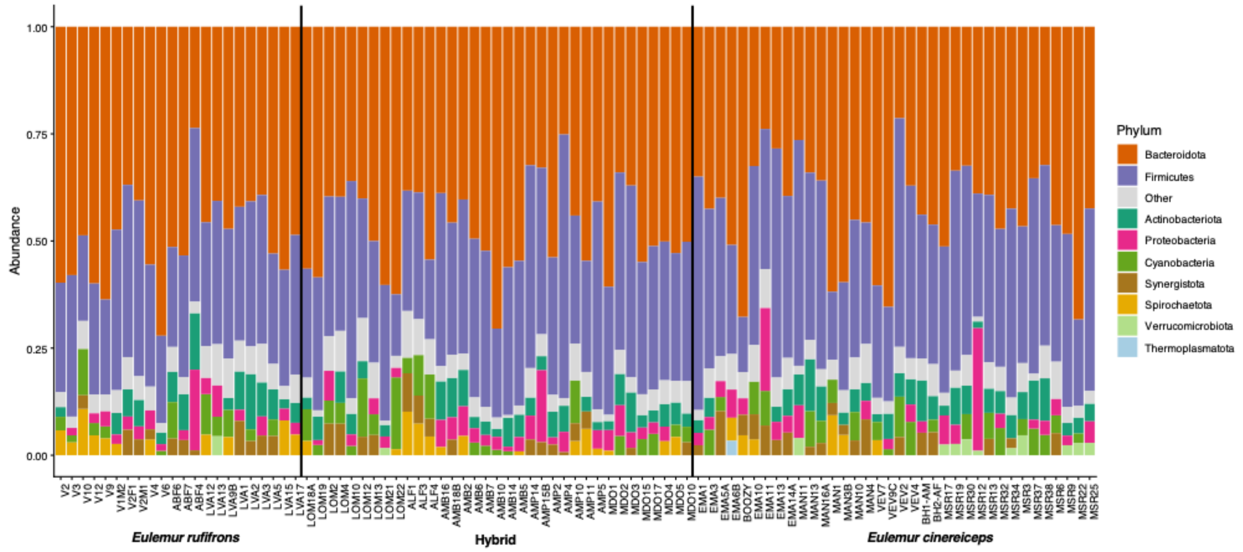


**Fig. S4:** Relative abundance of the dominant gut bacterial phyla. Each stacked bar represents a single sample, with phyla comprising the five most abundant taxa within each sample displayed individually and all remaining phyla combined into the "Other" category. Samples are grouped by host taxon, with vertical lines separating the three groups. Phyla are ordered in the legend according to their overall abundance across all samples.

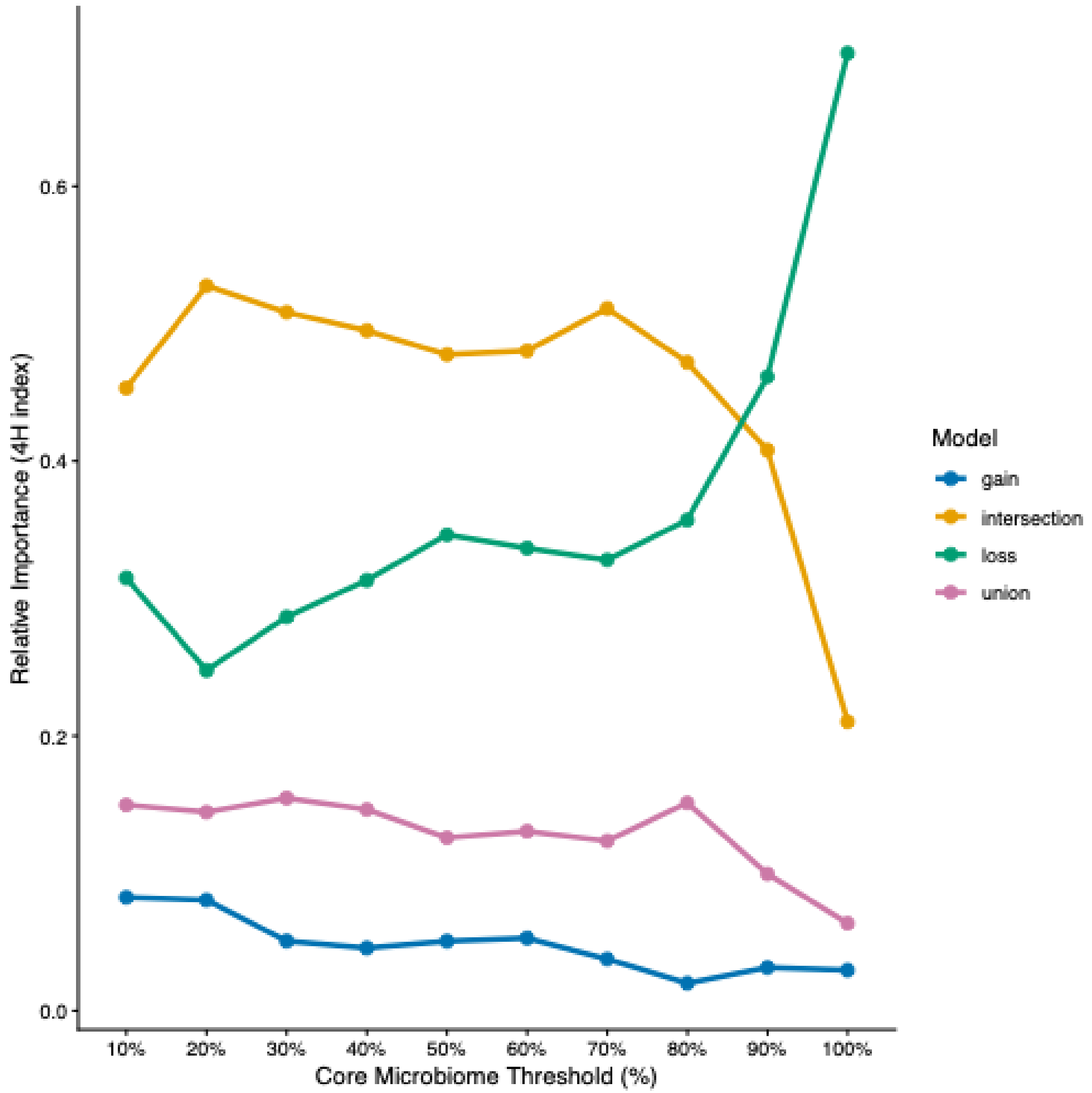


**Fig. S5:** Changes in the relative contribution of the four hybrid microbiome assembly models across core microbiome thresholds ranging from 10% to 100%. The 4H analysis partitions the hybrid gut microbiome into four components: intersection (microbial taxa shared between both parental species), gain (novel taxa unique to hybrids), loss (parental taxa absent from hybrids), and union (taxa present in either parental species but not shared). Values represent the centroid of 100 bootstrap replicates at each core microbiome threshold, with each bootstrap subsampled to the smallest host group size. Across most thresholds, the intersection component accounted for the largest proportion of the hybrid microbiome, whereas the gain and union components remained consistently low. The loss component increased with more stringent core thresholds.

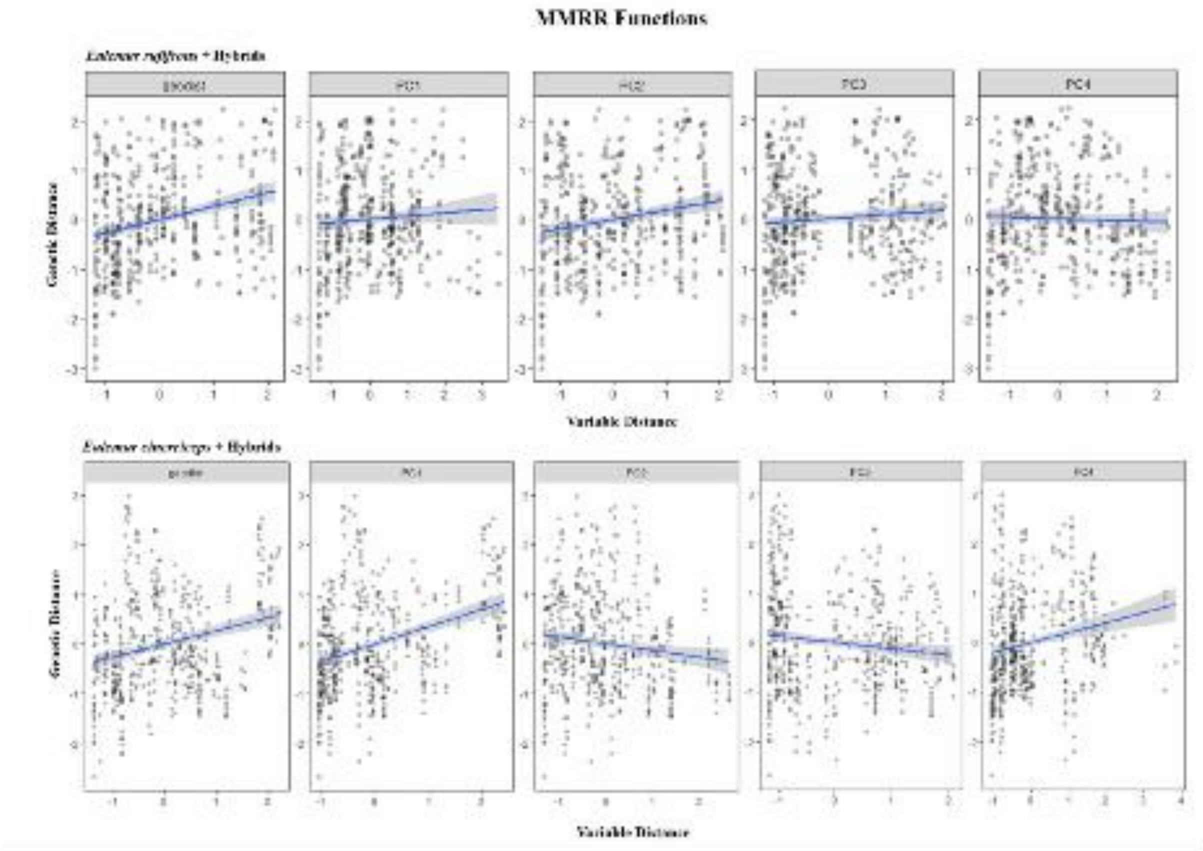


**Fig. S8:** Variable plots produced from MMRR analysis on Eulemur SNP data showing relationships between genetic distance and predictor variables. Plots in the upper panel (Eulemur rufifrons + hybrid samples) represent a weak linear relationship with geographic distance – indicating an IBD correlation, while plots in the lower panel (Eulemur cinereiceps + hybrid samples) represent a weak linear relationship with environmental variables from four PC-based BioClim environmental predictors – indicating an IBE correlation.

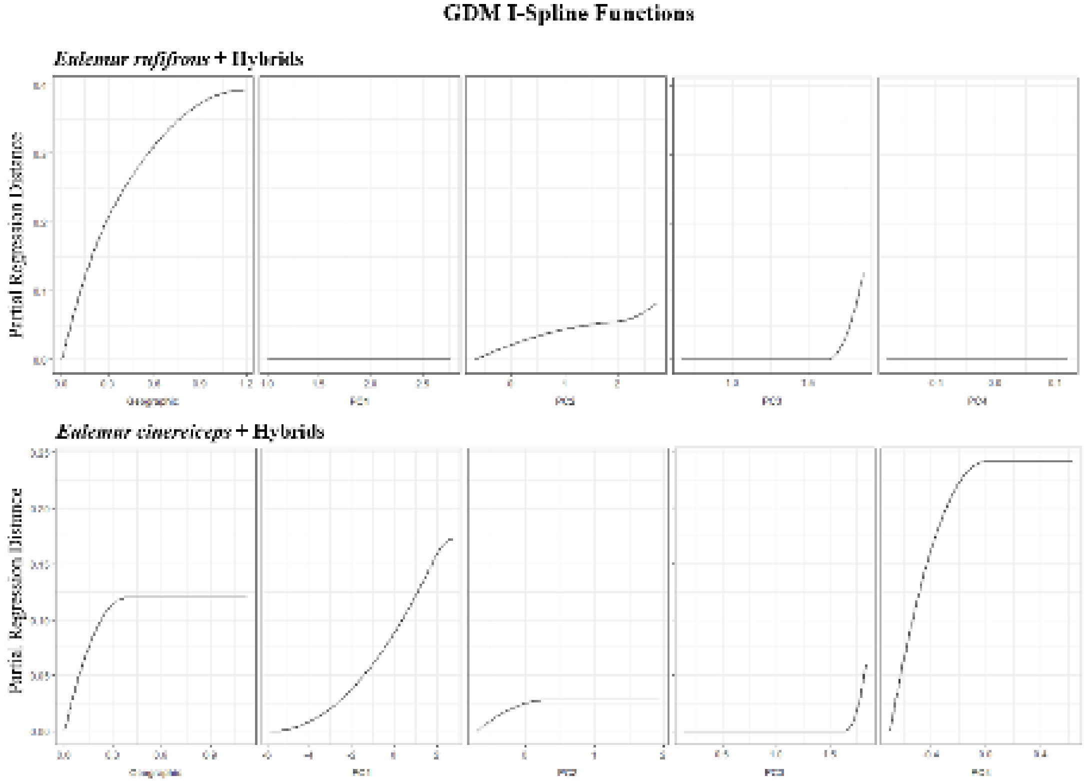


**Fig. S9:** I-spline function plots with fixed y-axes and free x-axes produced from fitted GDM analysis on Eulemur SNP data. Relative heights of the splines indicate the contribution of predictors to genetic distance dissimilarity across samples. For the environmental splines, PC1 represents all environmental variables and elevation, PC2 represents the effect of precipitation-centric BioClim variables, PC3 represents the effect of seasonality and annual temperature variation, and PC4 represents the effect of precipitation seasonality.

**Table S1:** Metadata for fecal samples used in population genomic analyses.

| Sample name | Species | Sex | Stage | Site | Latitude | Longitude | Collection date |
|---|---|---|---|---|---|---|---|
| ABF5 | *E. rufifrons* | Male | Adult | Ambatofisika | 21.85974 S | 47.3003 E | 1/27/22 |
| ABF6 | *E. rufifrons* | Female | Adult | Ambatofisika | 21.85974 S | 47.3003 E | 1/27/22 |
| ALF1 | Hybrid | Male | Adult | Ankona | 22.16197 S | 47.09542 E | 4/10/22 |
| ALF2 | Hybrid | Male | Adult | Ankona | 22.16197 S | 47.09542 E | 4/10/22 |
| ALF3 | Hybrid | Female | Adult | Ankona | 22.16197 S | 47.09542 E | 4/10/22 |
| ALF4 | Hybrid | Male | Adult | Ankona | 22.16197 S | 47.09542 E | 4/10/22 |
| AMB11 | Hybrid | Male | Adult | Ambarongy | 22.22994 S | 46.9952 E | 6/22/22 |
| AMB12 | Hybrid | Female | Juvenile | Ambarongy | 22.22994 S | 46.9952 E | 6/22/22 |
| AMB14 | Hybrid | Male | Adult | Ambarongy | 22.22994 S | 46.9952 E | 6/22/22 |
| AMB2 | Hybrid | Female | Adult | Ambarongy | 22.22171 S | 47.02002 E | 1/24/22 |
| AMP12 | Hybrid | Female | Juvenile | Ampasy | 22.30091 S | 47.02214 E | 6/27/22 |
| AMP15 | Hybrid | Female | Adult | Ampasy | 22.29395 S | 47.0019 E | 6/28/22 |
| AMP3 | Hybrid | Female | Adult | Ampasy | 22.29512 S | 47.00477 E | 1/31/22 |
| AMP6 | Hybrid | Male | Adult | Ampasy | 22.30161 S | 47.0216 E | 2/2/22 |
| AMP7 | Hybrid | Male | Adult | Ampasy | 22.30161 S | 47.0216 E | 2/2/22 |
| BH1 | *E. cinereiceps* | Male | Adult | Bemalaha | 22.96222 S | 47.17613 E | 6/1/22 |
| BH2 | *E. cinereiceps* | Male | Adult | Bemalaha | 22.96222 S | 47.17613 E | 6/1/22 |
| BOOZY | *E. cinereiceps* | Female | Juvenile | Emma | 22.41553 S | 47.17714 E | 7/27/22 |
| EMA5 | *E. cinereiceps* | Male | Adult | Emma | 22.41278 S | 47.17881 E | 5/27/22 |
| LOM22 | Hybrid | Female | Adult | Lomaka | 22.41278 S | 47.17881 E | 7/15/22 |
| LOM4 | Hybrid | Female | Adult | Lomaka | 22.04578 S | 47.17191 E | 4/2/22 |
| LOM7 | Hybrid | Female | Adult | Lomaka | 22.04578 S | 47.17191 E | 4/2/22 |
| LVA11 | *E. rufifrons* | Male | Juvenile | Lovasoa | 21.9315 S | 47.27229 E | 7/11/22 |
| LVA14 | *E. rufifrons* | Female | Adult | Lovasoa | 21.9315 S | 47.27229 E | 7/11/22 |
| LVA17 | *E. rufifrons* | Female | Adult | Lovasoa | 21.93312 S | 47.27633 E | 7/11/22 |
| LVA7 | *E. rufifrons* | Male | Adult | Lovasoa | 21.9355 S | 47.9355 E | 3/28/22 |

| | | | | | | | |
|---|---|---|---|---|---|---|---|
| LVA9 | *E. rufifrons* | Male | Adult | Lovasoa | 21.9315 S | 47.27229 E | 7/11/22 |
| MAN15 | *E. cinereiceps* | Male | Adult | Manasara | 22.61997 S | 47.21352 E | 8/1/22 |
| MAN2 | *E. cinereiceps* | Male | Adult | Manasara | 22.62687 S | 47.23049 E | 6/1/22 |
| MAN6 | *E. cinereiceps* | Male | Adult | Manasara | 22.63276 S | 47.24645 E | 6/4/22 |
| MAN8 | *E. cinereiceps* | Female | Adult | Manasara | 22.63276 S | 47.24645 E | 6/4/22 |
| MDO14 | Hybrid | Male | Adult | Mariandrano | 22.35533 S | 46.97911 E | 7/1/22 |
| MDO15 | Hybrid | Female | Adult | Mariandrano | 22.35533 S | 46.97911 E | 7/1/22 |
| MDO6 | Hybrid | Male | Adult | Mariandrano | 22.35637 S | 46.9718 E | 2/7/22 |
| MSR18 | *E. cinereiceps* | Male | Adult | Manombo | 23.01986 S | 47.72458 E | 3/3/20 |
| MSR26 | *E. cinereiceps* | Male | Adult | Manombo | 23.02061 S | 47.72422 E | 3/5/20 |
| MSR5 | *E. cinereiceps* | Female | Adult | Manombo | 23.02331 S | 47.72879 E | 2/27/20 |
| RNP25 | *E. rufifrons* | Male | Adult | Ranomafana | 21.26053 S | 47.42261 E | 2/11/20 |
| RNP36 | *E. rufifrons* | Female | Adult | Ranomafana | 21.26608 S | 47.42508 E | 7/20/20 |
| V11 | *E. rufifrons* | Male | Adult | Ranomafana | 21.2889 S | 47.43014 E | 3/5/22 |
| V12 | *E. rufifrons* | Male | Adult | Ranomafana | 21.2889 S | 47.43014 E | 3/5/22 |
| V8 | *E. rufifrons* | Female | Adult | Ranomafana | 21.2889 S | 47.43014 E | 3/5/22 |
| VEV5 | *E. cinereiceps* | Male | Adult | Vevembe | 22.79222 S | 47.19668 E | 6/10/22 |
| VEV7 | *E. cinereiceps* | Female | Adult | Vevembe | 22.79222 S | 47.19668 E | 6/12/22 |
| VEV8 | *E. cinereiceps* | Male | Adult | Vevembe | 22.79222 S | 47.19668 E | 6/12/22 |

**Table S2:** Metadata for fecal samples used in gut microbiome analyses.

| Sample name | Species | Sex | Stage | Site | Latitude | Longitude | Collection date |
|---|---|---|---|---|---|---|---|
| ABF4 | *E. rufifrons* | Male | Adult | Ambatofisika | 21.86142 S | 47.30562 E | 3/21/22 |
| ABF6* | *E. rufifrons* | Female | Adult | Ambatofisika | 21.85974 S | 47.30030 E | 3/22/22 |
| ABF7 | *E. rufifrons* | Male | Adult | Ambatofisika | 21.85974 S | 47.30030 E | 3/22/22 |
| ALF1* | Hybrid | Male | Adult | Ankona | 22.16197 S | 47.09542 E | 4/10/22 |
| ALF3* | Hybrid | Female | Adult | Ankona | 22.16197 S | 47.09542 E | 4/10/22 |
| ALF4* | Hybrid | Male | Adult | Ankona | 22.16197 S | 47.09542 E | 4/10/22 |
| AMB10 | Hybrid | Female | Adult | Ambarongy | 22.22994 S | 46.9952 E | 6/22/22 |
| AMB14* | Hybrid | Male | Adult | Ambarongy | 22.22994 S | 46.9952 E | 6/22/22 |
| AMB16 | Hybrid | Female | Adult | Ambarongy | 22.21662 S | 47.01621 E | 6/24/22 |
| AMB18 | Hybrid | Male | Adult | Ambarongy | 22.21662 S | 47.01621 E | 6/24/22 |
| AMB2 | Hybrid | Female | Adult | Ambarongy | 22.22171 S | 47.02002 E | 1/24/22 |
| AMB3 | Hybrid | Female | Juvenile | Ambarongy | 22.22171 S | 47.02002 E | 1/24/22 |
| AMB5 | Hybrid | Female | Adult | Ambarongy | 22.23038 S | 47.0053 E | 1/27/25 |
| AMB6 | Hybrid | Female | Adult | Ambarongy | 22.22950 S | 46.99849 E | 1/27/25 |
| AMB7 | Hybrid | Male | Adult | Ambarongy | 22.22950 S | 46.99849 E | 1/27/25 |
| AMP10 | Hybrid | Female | Adult | Ampasy | 22.30091 S | 47.02214 E | 6/27/25 |
| AMP11 | Hybrid | Male | Adult | Ampasy | 22.30091 S | 47.02214 E | 6/27/25 |
| AMP14 | Hybrid | Male | Adult | Ampasy | 22.29395 S | 47.00190 E | 6/28/25 |
| AMP15* | Hybrid | Female | Adult | Ampasy | 22.29395 S | 47.0019 E | 6/28/22 |
| AMP2 | Hybrid | Female | Adult | Ampasy | 22.29512 S | 47.00477 E | 1/30/22 |
| AMP4 | Hybrid | Male | Adult | Ampasy | 22.29512 S | 47.00477 E | 2/1/22 |
| AMP5 | Hybrid | Female | Adult | Ampasy | 22.30161 S | 47.02160 E | 2/2/22 |
| BH1* | *E. cinereiceps* | Male | Adult | Bemalaha | 22.96222 S | 47.17613 E | 6/1/22 |
| BH2* | *E. cinereiceps* | Male | Adult | Bemalaha | 22.96222 S | 47.17613 E | 6/1/22 |
| BOOZY* | *E. cinereiceps* | Female | Juvenile | Emma | 22.41553 S | 47.17714 E | 7/27/22 |
| EMA10 | *E. cinereiceps* | Female | Adult | Emma | 22.14145 S | 47.17436 E | 7/27/22 |

| | | | | | | | |
|---|---|---|---|---|---|---|---|
| EMA11 | *E. cinereiceps* | Male | Adult | Emma | 22.14145 S | 47.17436 E | 7/28/22 |
| EMA13 | *E. cinereiceps* | Male | Adult | Emma | 22.14145 S | 47.17436 E | 7/27/22 |
| EMA14 | *E. cinereiceps* | Female | Adult | Emma | 22.14145 S | 47.17436 E | 7/28/22 |
| EMA1 | *E. cinereiceps* | Male | Adult | Emma | 22.41435 S | 47.17442 E | 5/26/22 |
| EMA3 | *E. cinereiceps* | Female | Adult | Emma | 22.41435 S | 47.17442 E | 5/26/22 |
| EMA5 | *E. cinereiceps* | Male | Adult | Emma | 22.41435 S | 47.17442 E | 5/27/22 |
| EMA6 | *E. cinereiceps* | Female | Adult | Emma | 22.41435 S | 47.17442 E | 5/27/22 |
| LOM10 | Hybrid | Male | Adult | Lomaka | 22.06516 S | 47.16694 E | 4/3/22 |
| LOM12 | Hybrid | Male | Adult | Lomaka | 22.06516 S | 47.16694 E | 4/3/22 |
| LOM13 | Hybrid | Female | Adult | Lomaka | 22.06516 S | 47.16694 E | 4/3/22 |
| LOM18 | Hybrid | Female | Adult | Lomaka | 22.04337 S | 47.17248 E | 7/14/22 |
| LOM19 | Hybrid | Male | Adult | Lomaka | 22.04337 S | 47.17248 E | 7/14/22 |
| LOM21 | Hybrid | Male | Adult | Lomaka | 22.06536 S | 47.16697 E | 7/15/22 |
| LOM22* | Hybrid | Female | Adult | Lomaka | 22.41278 S | 47.17881 E | 7/15/22 |
| LOM2 | Hybrid | Male | Adult | Lomaka | 22.04578 S | 47.17191 E | 4/2/22 |
| LOM4* | Hybrid | Female | Adult | Lomaka | 22.04578 S | 47.17191 E | 4/2/22 |
| LVA12 | *E. rufifrons* | Female | Adult | Lovasoa | 21.93150 S | 47.27229 E | 7/11/22 |
| LVA13 | *E. rufifrons* | Male | Adult | Lovasoa | 21.93150 S | 47.27229 E | 7/11/22 |
| LVA15 | *E. rufifrons* | Male | Adult | Lovasoa | 21.93312 S | 47.27633 E | 7/11/22 |
| LVA17* | *E. rufifrons* | Female | Adult | Lovasoa | 21.93312 S | 47.27633 E | 7/11/22 |
| LVA1 | *E. rufifrons* | Male | Adult | Lovasoa | 21.93287 S | 47.27511 E | 3/27/22 |
| LVA2 | *E. rufifrons* | Male | Adult | Lovasoa | 21.93287 S | 47.27511 E | 3/27/22 |
| LVA3 | *E. rufifrons* | Female | Adult | Lovasoa | 21.93287 S | 47.27511 E | 3/27/22 |
| LVA5 | *E. rufifrons* | Female | Adult | Lovasoa | 21.27636 S | 47.27636 E | 3/28/22 |
| LVA9* | *E. rufifrons* | Male | Adult | Lovasoa | 21.9315 S | 47.27229 E | 7/11/22 |
| MAN10 | *E. cinereiceps* | Female | Adult | Manasara | 22.63276 S | 47.24645 E | 6/4/22 |
| MAN11 | *E. cinereiceps* | Male | Adult | Manasara | 22.61997 S | 47.21352 E | 8/1/22 |
| MAN13 | *E. cinereiceps* | Male | Adult | Manasara | 22.61997 S | 47.21352 E | 8/1/22 |

| | | | | | | | |
|---|---|---|---|---|---|---|---|
| MAN16 | *E. cinereiceps* | Female | Adult | Manasara | 22.61997 S | 47.21352 E | 8/1/22 |
| MAN1 | *E. cinereiceps* | Male | Adult | Manasara | 22.62687 S | 47.23049 E | 6/1/22 |
| MAN3 | *E. cinereiceps* | Female | Adult | Manasara | 22.62687 S | 47.23049 E | 6/1/22 |
| MAN4 | *E. cinereiceps* | Male | Adult | Manasara | 22.63276 S | 47.24645 E | 6/4/22 |
| MDO10 | *E. rufifrons* | Male | Adult | Mariandrano | 22.36007 S | 46.97812 E | 6/30/22 |
| MDO15* | *E. rufifrons* | Female | Adult | Mariandrano | 22.35533 S | 46.97911 E | 7/1/22 |
| MDO17 | *E. rufifrons* | Male | Adult | Mariandrano | 22.35533 S | 46.97911 E | 7/1/22 |
| MDO1 | *E. rufifrons* | Male | Adult | Mariandrano | 22.35357 S | 46.98151 E | 2/4/22 |
| MDO2 | *E. rufifrons* | Male | Adult | Mariandrano | 22.35357 S | 46.98151 E | 2/4/22 |
| MDO3 | *E. rufifrons* | Female | Adult | Mariandrano | 22.35357 S | 46.98151 E | 2/4/22 |
| MDO4 | *E. rufifrons* | Female | Adult | Mariandrano | 22.35637 S | 46.97180 E | 2/7/22 |
| MDO5 | *E. rufifrons* | Male | Adult | Mariandrano | 22.35637 S | 46.97180 E | 2/7/22 |
| MSR12 | *E. cinereiceps* | Female | Adult | Manombo | 23.0078 S | 47.70756 E | 3/2/20 |
| MSR13 | *E. cinereiceps* | Female | Adult | Manombo | 23.0078 S | 47.70756 E | 3/2/20 |
| MSR17 | *E. cinereiceps* | Male | Adult | Manombo | 23.0029 S | 47.71390 E | 3/3/20 |
| MSR19 | *E. cinereiceps* | Male | Adult | Manombo | 23.0029 S | 47.71390 E | 3/3/20 |
| MSR22 | *E. cinereiceps* | Male | Adult | Manombo | 23.02091 S | 47.72716 E | 3/4/20 |
| MSR25 | *E. cinereiceps* | Male | Adult | Manombo | 23.02371 S | 47.72430 E | 3/5/20 |
| MSR3 | *E. cinereiceps* | Male | Adult | Manombo | 23.01541 S | 47.72508 E | 2/21/20 |
| MSR30 | *E. cinereiceps* | Female | Adult | Manombo | 23.00315 S | 47.73323 E | 3/6/20 |
| MSR32 | *E. cinereiceps* | Female | Adult | Manombo | 23.01035 S | 47.73133 E | 3/11/20 |
| MSR34 | *E. cinereiceps* | Male | Adult | Manombo | 23.01035 S | 47.73133 E | 3/11/20 |
| MSR37 | *E. cinereiceps* | Male | Adult | Manombo | 23.01795 S | 47.72994 E | 3/13/20 |
| MSR38 | *E. cinereiceps* | Male | Adult | Manombo | 23.01795 S | 47.72994 E | 3/13/20 |
| MSR6 | *E. cinereiceps* | Male | Adult | Manombo | 23.02021 S | 47.71916 E | 2/28/20 |
| MSR9 | *E. cinereiceps* | Female | Adult | Manombo | 23.02021 S | 47.71916 E | 2/28/20 |
| V10 | *E. rufifrons* | Female | Adult | Ranomafana | 21.2889 S | 47.43014 E | 3/5/22 |
| V12 | *E. rufifrons* | Male | Adult | Ranomafana | 21.2889 S | 47.43014 E | 3/5/22 |

| | | | | | | | |
|---|---|---|---|---|---|---|---|
| V1M2 | *E. rufifrons* | Male | Adult | Ranomafana | 21.28904 S | 47.42889 E | 7/20/22 |
| V2F1 | *E. rufifrons* | Female | Adult | Ranomafana | 21.28904 S | 47.42889 E | 7/20/22 |
| V2M1 | *E. rufifrons* | Male | Adult | Ranomafana | 21.28904 S | 47.42889 E | 7/20/22 |
| V2 | *E. rufifrons* | Male | Adult | Ranomafana | 21.2881 S | 47.42970 E | 1/15/22 |
| V3 | *E. rufifrons* | Female | Adult | Ranomafana | 21.2881 S | 47.42970 E | 1/15/22 |
| V4 | *E. rufifrons* | Male | Adult | Ranomafana | 21.29138 S | 47.43319 E | 1/16/22 |
| V6 | *E. rufifrons* | Female | Adult | Ranomafana | 21.29138 S | 47.43319 E | 1/16/22 |
| V9 | *E. rufifrons* | Male | Adult | Ranomafana | 21.2889 S | 47.43014 E | 3/5/22 |
| VEV2 | *E. cinereiceps* | Female | Adult | Vevembe | 22.79782 S | 47.18822 E | 6/10/22 |
| VEV4 | *E. cinereiceps* | Male | Adult | Vevembe | 22.79782 S | 47.18822 E | 6/10/22 |
| VEV7* | *E. cinereiceps* | Female | Adult | Vevembe | 22.79222 S | 47.19668 E | 6/12/22 |
| VEV9 | *E. cinereiceps* | Male | Adult | Vevembe | 22.79222 S | 47.19668 E | 6/12/22 |

**Table S3**: Pairwise weighted FST values between *E. cinereiceps*, *E. rufifrons*, and putative hybrids

| | *E. cinereiceps* | *E. rufifrons* | Hybrid |
|---|---|---|---|
| *E. cinereiceps* | - | | |
| *E. rufifrons* | 0.117982 | - | |
| Hybrid | 0.092079 | 0.024268 | - |

**Table S4:** Results of MMRR analyses for IBD and IBE on *Eulemur* samples. Models were run using SNP data from samples binned into three comparative test groups: (A) *Eulemur rufifrons* and hybrids, (B) *Eulemur cinereiceps* and hybrids, and (C) all samples treated as a single deme. Variables contributing the most explanatory variance to the model are bolded and indicated with an asterisk (*). PCs 1-4 explain the effect of environmental variables, while the "GeoDist" variable was based on geographic distance matrices derived from coordinates of the *Eulemur* samples collected in this study.

| **A. *Eulemur rufifrons* + Hybrid Model** | | | | |
|---|---|---|---|---|
| **Variable** | **Estimate** | **p** | **95% Lower** | **95% Upper** |
| GeoDist | **0.28*** | 0.150 | 0.160 | 0.390 |
| Intercept | 0.000 | 0.300 | -0.080 | 0.080 |
| PC1 | -0.060 | 0.550 | -0.150 | 0.030 |
| PC2 | 0.070 | 0.570 | -0.020 | 0.170 |
| PC3 | -0.010 | 0.920 | -0.110 | 0.080 |
| PC4 | -0.050 | 0.500 | -0.130 | 0.040 |
| **R-Squared** | **0.090** | - | - | - |
| **F-Statistic** | **8.030** | - | - | - |
| **F p-value** | **0.090** | - | - | - |
| **B. *Eulemur cinereiceps* + Hybrid Model** | | | | |
| **Variable** | **Estimate** | **p** | **95% Lower** | **95% Upper** |
| GeoDist | -0.15 | 0.44 | -0.32 | 0.02 |
| Intercept | 0 | 0.07 | -0.07 | 0.07 |
| PC1 | **0.6*** | 0.002 | 0.39 | 0.81 |
| PC2 | -0.14 | 0.18 | -0.21 | -0.07 |
| PC3 | -0.13 | 0.08 | -0.21 | -0.05 |
| PC4 | -0.15 | 0.21 | -0.27 | -0.04 |
| **R-Squared** | **0.17** | - | - | - |
| **F-Statistic** | **20.78** | - | - | - |
| **F p-value** | **0.01** | - | - | - |
| **C. *Eulemur* All Samples Model** | | | | |

| Variable | Estimate | p | 95% Lower | 95% Upper |
|---|---|---|---|---|
| GeoDist | **0.23** | 0.1 | 4.33E-16 | 0.3 |
| Intercept | 0.00 | 0.19 | 0.15 | 0.05 |
| PC1 | **0.29*** | 0.05 | -0.05 | 0.38 |
| PC2 | 0.07 | 0.37 | 0.2 | 0.13 |
| PC3 | -0.17 | 0.01 | 0.01 | -0.12 |
| PC4 | -0.07 | 0.57 | -0.22 | 0.003 |
| **R-Squared** | **0.186** | - | - | - |
| **F-Statistic** | **43.02** | - | - | - |
| **F p-value** | **0.004** | - | - | - |

**Table S5:** Results of GDM analyses for IBD and IBE on *Eulemur* samples. Models were run using SNP data from samples binned into three comparative test groups: (A) *Eulemur rufifrons* and hybrids, (B) *Eulemur cinereiceps* and hybrids, and (C) all samples treated as a single deme. Variables contributing the most null deviance to the model are bolded and indicated with an asterisk (*). PCs 1-4 explain the effect of environmental variables, while the "GeoDist" variable was based on geographic distance matrices derived from coordinates of the *Eulemur* samples collected in this study.

| **A. *Eulemur rufifrons* + Hybrid Model** | | | |
|---|---|---|---|
| **Predictor** | **Coefficient** | **Predictor Importance** | **Predictor p-value** |
| GeoDist | 0.392 | 37.81* | 0.001 |
| PC1 | 0.000 | n/a | n/a |
| PC2 | 0.080 | 15.240 | 0.880 |
| PC3 | 0.130 | 11.830 | 0.730 |
| PC4 | 0.000 | n/a | n/a |
| **Model Deviance** | 26.570 | - | - |
| **% Deviance Explained** | 9.690 | - | - |
| **Model p-value** | 0.360 | - | - |
| **Fitted Permutations** | 1000 | - | - |
| **B. *Eulemur cinereiceps* + Hybrid Model** | | | |

| Predictor | Coefficient | Predictor Importance | Predictor p-value |
|---|---|---|---|
| GeoDist | 0.12 | 4.71 | 0.15 |
| PC1 | 0.17 | 6.3 | 0.66 |
| PC2 | 0.03 | **14.19*** | 0.79 |
| PC3 | 0.06 | 9.98 | 0.74 |
| PC4 | 0.24 | 7.01 | 0.64 |
| **Model Deviance** | 30.130 | - | - |
| **% Deviance Explained** | 13.900 | - | - |
| **Model p-value** | 0.500 | - | - |
| **Fitted Permutations** | 1000 | - | - |
| **C. *Eulemur* All Samples Model** | | | |
| **Predictor** | **Coefficient** | **Predictor Importance** | **Predictor p-value** |
| GeoDist | 0.23 | 4.58 | 0.03 |
| PC1 | 0.18 | 7.13 | 0.55 |
| PC2 | 0.13 | 7.4 | 0.55 |
| PC3 | 0.04 | 5.05 | 0.78 |
| PC4 | 0.17 | 3.04 | 0.66 |
| **Model Deviance** | 42.81 | - | - |
| **% Deviance Explained** | 17.82 | - | - |
| **Model p-value** | 0.17 | - | - |
| **Fitted Permutations** | 1000 | - | - |

**Table S6**: Results of permutational multivariate analysis of variance (PERMANOVA) conducted in QIIME 2 using the adonis function (999 permutations) on four beta diversity distance metrics (Bray–Curtis, Jaccard, unweighted UniFrac, and weighted UniFrac). $R^2$ values represent the proportion of variation explained by each factor, and p-values indicate significance based on permutation tests. Significant effects were observed for Species, Site, and the Site × Group interaction across most distance metrics, whereas Sex was not a significant predictor of community variation. Residuals represent unexplained variation.

| Metric | Variable | R2 | P-value |
|---|---|---|---|
| Bray-Curtis | Species | 0.09 | 0.001 |
| | Site | 0.20 | 0.001 |
| | Sex | 0.01 | 0.603 |
| | Site:Group | 0.14 | 0.001 |
| | Residuals | 0.57 | N/A |
| Jaccard | Species | 0.08 | 0.001 |
| | Site | 0.19 | 0.001 |
| | Sex | 0.01 | 0.749 |
| | Site:Group | 0.12 | 0.001 |
| | Residuals | 0.60 | N/A |
| Unweighted UniFrac | Species | 0.04 | 0.001 |
| | Site | 0.18 | 0.001 |
| | Sex | 0.01 | 0.154 |
| | Site:Group | 0.14 | 0.005 |
| | Residuals | 0.63 | N/A |
| Weighted UniFrac | Species | 0.09 | 0.001 |
| | Site | 0.18 | 0.004 |
| | Sex | 0.01 | 0.242 |
| | Site:Group | 0.18 | 0.001 |
| | Residuals | 0.53 | N/A |

**Table S7**: Results of Mantel tests examining correlations between beta diversity distance matrices (Bray–Curtis, Jaccard, unweighted UniFrac, and weighted UniFrac) and explanatory distance matrices (genetic, geographic, and environmental). Analyses were conducted using Spearman's rank correlation with 999 permutations. Reported values represent Mantel correlation coefficients (r) and associated p-values. Across metrics, beta diversity showed consistent and significant correlations with genetic distance, while relationships with geographic and environmental distance were weaker, variable among metrics, and often not statistically significant.

| Metric | Formula | R2 | P-Value |
|---|---|---|---|
| Bray-Curtis | xdis = Beta diversity<br>ydis = genetic | 0.25 | 0.024 |
| | xdis = Beta diversity<br>ydis = geographic | 0.05 | 0.285 |
| | xdis = Beta diversity<br>ydis = environment | 0.10 | 0.154 |
| Jaccard | xdis = Beta diversity<br>ydis = genetic | 0.26 | 0.013 |
| | xdis = Beta diversity<br>ydis = geographic | 0.24 | 0.006 |
| | xdis = Beta diversity | 0.19 | 0.043 |

| | | | |
|---|---|---|---|
| | ydis = environment | | |
| Unweighted UniFrac | xdis = Beta diversity<br>ydis = genetic | 0.26 | 0.020 |
| | xdis = Beta diversity<br>ydis = geographic | 0.04 | 0.303 |
| | xdis = Beta diversity<br>ydis = environment | 0.05 | 0.300 |
| Weighted UniFrac | xdis = Beta diversity<br>ydis = genetic | 0.22 | 0.038 |
| | xdis = Beta diversity<br>ydis = geographic | -0.06 | 0.726 |
| | xdis = Beta diversity<br>ydis = environment | 0.00 | 0.433 |

**Table S8**: Results of Partial Mantel tests assessing the relationships between beta diversity (Bray–Curtis, Jaccard, unweighted UniFrac, and weighted UniFrac distance matrices) and explanatory distance matrices (genetic, geographic, and environmental), while controlling for the effect of a third matrix. Analyses were conducted using Spearman's rank correlation with 999 permutations. Reported values represent Mantel correlation coefficients (r) and associated p-values. Significant correlations were observed between beta diversity and genetic distance after controlling for geographic or environmental distance, whereas associations with environmental or geographic distance alone were generally weak and non-significant.

| Metric | Formula | R2 | P-Value |
|---|---|---|---|
| Bray-curtis | xdis= Beta diversity<br>ydis= genetic<br>zdis = geographic | 0.25 | 0.016 |
| | xdis= Beta diversity<br>ydis = genetic<br>zdis = environment | 0.23 | 0.031 |
|  | xdis = Beta diversity<br>ydis = environment<br>zdis = geographic | 0.08 | 0.221 |
|  | xdis = Beta diversity<br>ydis = geographic<br>zdis = environment | 0.00 | 0.472 |
| Jaccard | xdis= Beta diversity<br>ydis= genetic<br>zdis = geographic | 0.19 | 0.049 |
| | xdis= Beta diversity<br>ydis = genetic<br>zdis = environment | 0.22 | 0.033 |
|  | xdis = Beta diversity<br>ydis = environment<br>zdis = geographic | 0.08 | 0.214 |
|  | xdis = Beta diversity<br>ydis = geographic<br>zdis = environment | 0.17 | 0.048 |
| Unweighted UniFrac | xdis= Beta diversity<br>ydis= genetic<br>zdis = geographic | 0.27 | 0.021 |
| | xdis= Beta diversity<br>ydis = genetic | 0.26 | 0.011 |

| | | | |
|---|---|---|---|
| | zdis = environment | | |
| | xdis = Beta diversity<br>ydis = environment<br>zdis = geographic | 0.03 | 0.378 |
| | xdis = Beta diversity<br>ydis = geographic<br>zdis = environment | 0.02 | 0.382 |
| Weighted UniFrac | xdis= Beta diversity<br>ydis= genetic<br>zdis = geographic | 0.26 | 0.015 |
| | xdis= Beta diversity<br>ydis = genetic<br>zdis = environment | 0.226 | 0.018 |
| | xdis = Beta diversity<br>ydis = environment<br>zdis = geographic | 0.04 | 0.365 |
| | xdis = Beta diversity<br>ydis = geographic<br>zdis = environment | -0.07 | 0.755 |

**Table S9:** Summary of MaAsLin2 results comparing KO abundances between hybrids and parental species. The "KEGG Level 1" and "KEGG Level 2" columns denote functional pathway hierarchies based on BRITE annotations. "Comparison" specifies which parental species differs significantly from hybrids. For each Level 2 category, the "Average log-transformed coefficient" represents the mean effect size across all significantly associated KOs, where positive values indicate enrichment in the parental species relative to hybrids. "# of KOs" indicates the total number of significant KOs contributing to each functional category–comparison pair.

| KO Level 1 | KO Level 2 | Comparison | Average log-transformed coefficient | # of KOs |
|---|---|---|---|---|
| Metabolism | Amino acid metabolism | *E. cinereiceps* | 0.55 | 15 |
| Metabolism | Carbohydrate metabolism | *E. cinereiceps* | 0.36 | 19 |
| Metabolism | Energy metabolism | *E. rufifrons* | 1.63 | 1 |
| Metabolism | Energy metabolism | *E. cinereiceps* | 0.78 | 6 |
| Metabolism | Fructose and mannose metabolism | *E. cinereiceps* | -2.08 | 1 |
| Metabolism | Glycan biosynthesis and metabolism | *E. cinereiceps* | 0.64 | 11 |
| Metabolism | Glycerophospholipid metabolism | *E. cinereiceps* | 0.37 | 1 |
| Metabolism | Lipid metabolism | *E. cinereiceps* | -0.26 | 6 |
| Metabolism | Metabolism of cofactors and vitamins | *E. cinereiceps* | 0.38 | 13 |
| Metabolism | Methane metabolism | *E. cinereiceps* | 0.54 | 1 |
| Metabolism | Nucleotide metabolism | *E. cinereiceps* | 0.43 | 11 |
| Metabolism | Propanoate metabolism | *E. cinereiceps* | -1.73 | 2 |

| | | | | |
|---|---|---|---|---|
| Metabolism | Protein families: metabolism | *E. cinereiceps* | 0.25 | 17 |
| Metabolism | Pyrimidine metabolism | *E. cinereiceps* | -0.1 | 1 |
| Metabolism | Sulfur metabolism | *E. cinereiceps* | 0.87 | 2 |
| Metabolism | Unclassified: metabolism | *E. cinereiceps* | 0.16 | 17 |
| Cellular processes | Cell growth and death | *E. cinereiceps* | -0.07 | 2 |
| Cellular processes | Cell motility | *E. cinereiceps* | 0.64 | 1 |
| Cellular processes | Quorum sensing | *E. cinereiceps* | 0.31 | 3 |
| Protein families: signaling and cellular processes | Antimicrobial resistance genes | *E. cinereiceps* | 0.87 | 4 |
| Protein families: signaling and cellular processes | Bacterial motility proteins | *E. cinereiceps* | 0.79 | 1 |
| Protein families: signaling and cellular processes | Prokaryotic defense system | *E. cinereiceps* | 0.77 | 8 |
| Protein families: signaling and cellular processes | Transporters | *E. cinereiceps* | 0.41 | 14 |
| Unclassified: signaling and cellular processes | Others: putative iron-regulated protein | *E. cinereiceps* | 0.74 | 1 |
| Unclassified: signaling and cellular processes | Signaling proteins | *E. cinereiceps* | 0.14 | 3 |
| Unclassified: signaling and cellular processes | Structural proteins | *E. cinereiceps* | 0.32 | 1 |
| Unclassified: signaling and cellular processes | Transport | *E. cinereiceps* | 0.06 | 2 |
| Environmental Information Processing | Membrane transport | *E. cinereiceps* | 0.27 | 19 |
| Environmental Information Processing | Signal transduction | *E. rufifrons* | 2.1 | 2 |
| Environmental Information Processing | Signal transduction | *E. cinereiceps* | 0.26 | 8 |
| Genetic Information Processing | Folding, sorting and degradation | *E. cinereiceps* | 0.06 | 11 |
| Genetic Information Processing | Replication and repair | *E. cinereiceps* | -0.04 | 16 |
| Genetic Information Processing | Transcription | *E. cinereiceps* | -0.04 | 1 |
| Genetic Information Processing | Translation | *E. cinereiceps* | -0.04 | 53 |
| Protein families: genetic information processing | Chaperones and folding catalysts | *E. cinereiceps* | -0.44 | 5 |

| | | | | |
|---|---|---|---|---|
| Protein families: genetic information processing | Chromosome and associated proteins | *E. cinereiceps* | -0.15 | 3 |
| Protein families: genetic information processing | DNA repair and recombination proteins | *E. cinereiceps* | 0.91 | 5 |
| Protein families: genetic information processing | DNA replication proteins | *E. cinereiceps* | -0.03 | 1 |
| Protein families: genetic information processing | Ribosome biogenesis | *E. cinereiceps* | -0.04 | 13 |
| Protein families: genetic information processing | Transcription factors | *E. rufifrons* | -1.2 | 1 |
| Protein families: genetic information processing | Transcription factors | *E. cinereiceps* | 0.06 | 11 |
| Protein families: genetic information processing | Transcription machinery | *E. cinereiceps* | -0.04 | 2 |
| Protein families: genetic information processing | Transfer RNA biogenesis | *E. cinereiceps* | 0.24 | 6 |
| Protein families: genetic information processing | Translation factors | *E. cinereiceps* | -0.04 | 11 |
| Unclassified: genetic information processing | Replication and repair | *E. cinereiceps* | -0.55 | 2 |
| Unclassified: genetic information processing | Protein processing | *E. cinereiceps* | -0.04 | 1 |
| Organismal Systems | Endocrine system | *E. cinereiceps* | 1.25 | 1 |
| Organismal Systems | Immune system | *E. cinereiceps* | 0.06 | 1 |
| Organismal Systems | Sensory system | *E. cinereiceps* | 0.73 | 1 |
