## supplemental for "Gene flow erodes core gut microbiota, but not microbiome function, across a natural primate hybrid zone"

### SUPPLEMENTAL MATERIALS

#### Methods

qPCR to confirm presence of mammalian DNA: We randomly selected 10 pooled FecalSeq eluates for qPCR to verify we had achieved detectable amounts of mammalian DNA. We amplified the mammalian MYCBP gene using *cmyc* primers (Morin et al. 2001). qPCR cocktails contained 20 µl of 1X SYBR Green master mix, 1 µl of each primer (forward and reverse), and 1 µl of DNA. We used a positive control with 55 ng/µl of starting DNA from a lemur (*Indri indri*) to estimate our mammalian DNA yields. Amplification was then performed using a qPCR instrument as follows: 95°C for 15 minutes, followed by 50 cycles of 94°C for 15s, 59°C for 25s, and 72°C for 20s.

Fecal-seq specifications: Laboratory procedures closely followed the supplemental protocol outlined in Chiou & Bergey (2018) with the following specifications/exceptions: (1) we prepared all MBD-Fc-bound magnetic beads in 160 µl batches; (2) we used 30 µl of starting DNA; (3) we always performed the "optional" second wash (step 24); (4) we tripled the amount of time spent mixing the beads with elution buffer (2M NaCl) in step 29; and (5) we used AMPure XP Bead-Based Reagent instead of a homebrew for the Auxiliary A bead cleanup protocol. As an additional modification to maximize total DNA yield, we performed the entire FecalSeq procedure three times per sample and pooled the final eluates.

Microbiome PCR: PCRs contained 22.5 µL of Accuprime Pfx Supermix™, 4 µL of DNA, and 1 µL each of forward and reverse primer (10 mM concentration). PCR amplification was performed in a thermal cycler with a touchdown protocol as follows: a 2min initial denaturation at 95°C followed by 19 cycles of 20s at 95°C, 15s at 60°C, and 5min at 72°C, an additional 19 cycles of 20s at 95°C, 15s of 55°C, and 5min at 72°C, and then a final extension of 10min at 72°C. Size and quality of PCR products were confirmed using agarose gel electrophoresis. To confirm adapters had ligated, we randomly selected 10 PCRs for qPCR quantification using the NEBNext® Library Quant Kit for Illumina®. qPCR cocktails contained 16 µL of NEBNext® Library Quant Master Mix (with primers) and 4 µL of either DNA standards or library (PCR product) dilution. In preparation for qPCR, libraries were diluted 1:1,000, 1:10,000, and 1:100,000. We then ran the qPCR with an initial denaturation of 95°C for 1 minute followed by 35 cycles of denaturation at 95°C for 15s and extension at 63°C for 45s.

Bioinformatics: Low coverage lemur genomic data**:** Quality assessment, quality control, and processing of sequence data was accomplished using the nf-core pipeline sarek (Garcia et al., 2020). Adapters and low-quality bases were removed using fastp (Chen et al., 2018). Filtered reads were mapped to the *E. rufifrons* reference genome (GCF_041146395.1) with bwa-mem (Li and Durbin, 2019) and further processed using gatk v4.1.4.1 (De Summa et al., 2017). The quality filtered binary alignment files were then used as input for angsd v.0.929.55 (Korneliussen et al., 2014). The samtools model (-GL 1) was specified and only bases with a phred score ≥ 20 (-minQ 20) and a mapping quality ≥ 20 (-minMapQ 20) were retained. To ensure sites were well represented among individuals, only biallelic sites in at least 80% of the samples (n=36) with a minor allele frequency of 0.025 and a *p-value* cutoff of $10^{-6}$ were used for analyses. As some of our downstream analyses required called genotypes or site frequency spectra rather than genotype likelihoods, we also ran angsd using the "-doGeno" and "-doSaf" flags with identical filters.

Landscape genomic input data: The framework for both the MMRR and GDM analyses used the same suite of input data. Coordinate data for each lemur fecal sample was collected in the field and collated into pairwise geographic distance matrices that corresponded to respective focal species-hybrid comparison groups. We downloaded 30-arc second (1km) raster layers for 19 Bioclimatic variables for our study area in southeast Madagascar, as well as a corresponding 30 arc-second NASA STRM digital elevation model (DEM) raster of the same bounding region produced from an open-source shapefile obtained from HDX v1.82.10 (https://data.humdata.org) using QGIS 3.30.2. We performed raster PCA on the resultant stack of input layers in R using packages *terra* and *RStoolbox* to mitigate spatial collinearity between environmental data layers and lemur sampling localities and subsequently extracted the final environmental input data from layers comprising 98% of the total raster PCA model variance. The final inputs for MMRR and GDM were (per-scenario) genetic distance matrices calculated from our filtered SNP data using PLINK 1.9 (Purcell et al. 2007).

For the MMRR analysis, geographic and environmental distances served as predictor variable regression coefficients (β) in the MMRR, while genetic distances served as response variables. The incorporation of the randomization procedure inherent to MMRR mitigated statistical non-independence of values within both predictor matrices (Wang 2013). The MMRR models were run for 1,000 permutations and fitted with the incorporation of a backwards model selection procedure to obtain the final geographic- and environmental PC-genetic distance correlations. Similarly, GDM runs utilized geographic- and BioClim raster PC-derived environmental distances as predictors, while the PLINK distance matrix values served as the response. The GDM ran for 1,000 permutations and site-pair comparisons of compositional dissimilarity were gauged for importance via summary statistics and the relative heights of the resultant I-spline functions.

Base scripts and associated package dependency descriptions are available at https://github.com/TheWangLab/algatr.

**Results**

Landscape genomics: The orthogonal function loadings from the RasterPCA model for our study area resulted in the first four environmental PCS explaining 98.35% of the total model variance. PC1 was described by the combination of all environmental BioClim variables and elevation, comprising 79.39% of the cumulative model variance. PC2 was described mostly by temperature related BioClim variables, comprising 13.87% of the cumulative model variance. PC3 was described by precipitation related BioClim variables, comprising 3.46% of the cumulative model variance. PC4 was primarily described by precipitation related BioClim variables, comprising 1.62% of cumulative model variance.

For the *E. rufifrons*-hybrid group (ERH), MMRR best fit model selection resulted in a reduced model solely comprised of environmental PCs (PC2 + PC4) indicating a statistically significant IBE correlation with genetic distance ($\beta_{PC2} = 0.20$, $P \leq 0.01$; $\beta_{PC4} = 0.14$, $P \leq 0.01$) (Fig. S8; Table S4). For the *E. cinereiceps*-hybrid group (ECH), MMRR analysis revealed no best fit model with variable reduction and the full model including all environmental PCs and geographic distance was not significant ($\beta_{PC1} = 0.19$, $P \leq 0.31$; $\beta_{PC2} = -0.07$, $P \leq 0.58$; $\beta_{PC3} = -0.11$, $P \leq 0.33$; $\beta_{PC4} = -0.04$, $P \leq 0.80$; $\beta_{GeoDist} = 0.17$, $P \leq 0.81$) (Fig. S8; Table S4). For the all samples group (AS), best fit model selection yielded a reduced model solely comprised of geographic distance as the predictor, indicating a signal of IBD ($\beta_{GeoDist} = 0.29$, $P \leq 0.01$) (Fig. S8; Table S4).

GDM model analyses for ERH resulted in a best fit model with PC3 removed; however, the effects of environmental or geographic distance on genetic compositional dissimilarity were not highly supported – 12.60% of overall deviance was explained by the non-significant model ($P \leq 0.06$) (Fig. S9; Table S5). PC1 and PC2 explained the majority of overall deviance, indicating environmental variables were the strongest predictors within the (weakly supported) model (Fig. S9; Table S5). GDM results for ECH samples yielded no reduced model and the strongest predictors within the model were PC2, PC3, and PC4 (Table S5). The GDM had low overall percent deviance explained by the model (13.64%) and it was non-significant for any individual predictor (Table S5). For AS, the best fit model included geographic distance, PC1, PC2, and PC4 as predictors, though again the GDM had low overall percent deviance explained by the model (12.07%) and a non-significant result was recovered (Fig. S9; Table S5). Overall, IBD-IBE analyses only detected a weak signal of IBE between the ERH samples, and no significant effects of IBD on population genetic variation were detected in any of the *Eulemur* sample groups.
